# Tight junction ZO proteins maintain tissue fluidity, ensuring efficient collective cell migration

**DOI:** 10.1101/2021.03.10.434539

**Authors:** Mark Skamrahl, Hongtao Pang, Maximilian Ferle, Jannis Gottwald, Angela Rübeling, Riccardo Maraspini, Alf Honigmann, Tabea A. Oswald, Andreas Janshoff

**Affiliations:** University of Göttingen, Institute of Physical Chemistry, Tammannstr. 6, 37077 Göttingen, Germany; University of Göttingen, Institute of Organic and Biomolecular Chemistry, Tammannstr. 2, 37077 Göttingen, Germany; Max Planck Institute of Molecular Cell Biology and Genetics, Pfotenhauerstraße 108, 01307 Dresden, Germany

**Keywords:** tight junctions, collective cell migration, jamming, cell mechanics, atomic force microscopy

## Abstract

Tight junctions are essential components of epithelial tissues connecting neighboring cells to provide protective barriers. Albeit their general function to seal compartments is well understood, their role in collective cell migration is largely unexplored. Here, the importance of the tight junction proteins ZO1 and ZO2 for epithelial migration is investigated employing video microscopy in conjunction with velocimetry, segmentation, cell tracking, and atomic force microscopy/spectroscopy. The results indicate that ZO proteins are necessary for fast and coherent migration. In particular, ZO1 and 2 loss (dKD) induces actomyosin remodeling away from the central cortex towards the periphery of individual cells, resulting in altered viscoelastic properties. A tug-of-war emerges between two subpopulations of cells with distinct morphological and mechanical properties: 1) smaller and highly contractile cells with an outward-bulged apical membrane, and 2) larger, flattened cells, which, due to tensile stress, display a higher proliferation rate. In response, the cell density increases, leading to crowding-induced jamming and more small cells over time. Co-cultures comprising wildtype and dKD cells display phase separation based on differences in contractility rather than differential adhesion. This study shows that ZO proteins are necessary for efficient collective cell migration by maintaining tissue fluidity and controlling proliferation.

## 1. Introduction

Cellular junctions endow epithelial tissues with their barrier functions by physically connecting neighboring cells. Junction integrity is critical to prevent many diseases. While, among the various junction types, adherens junctions are typically considered as mechanical couplers between cells in epithelia, recent evidence also suggests an important mechanical role for tight junctions (TJs).^[1–7]^ It is conceived that TJs provide a mechanical feedback system regulating the contractility of individual cells via the actomyosin cytoskeleton and their adhesion strength to neighboring cells.^[1,4,8–11]^ Specifically, it was shown that TJs provide a negative mechanical feedback to individual cells in a layer, so that they contract less, lowering the forces on the adherens junctions.^[1,4]^ Once TJ formation is inhibited, cells respond by building thick actomyosin rings at the cell periphery, which, upon contraction, lead to severe heterogeneity of the cell morphology, particularly visible at the apical side.^[8,4,12,11]^ Since this mechanical TJ-based mechanism was established only recently, explicit knowledge of its implications for crucial biological processes such as collective migration remains limited. Collective cell migration depends on an intricate interplay of the mechanical interaction in a cell layer ranging from single cells, e.g., leader cells at the advancing migration front, to the collective behavior of the cell sheet on a mesoscopic level.^[13–19]^ This interplay depends on the fine tuning of cell motility, density, contractility, and cell-cell adhesion.^[20–32]^

Important advances have been achieved in understanding how collective cell migration is generally influenced by the adhesion-mediating junction proteins.^[28]^ However, there is controversial evidence on the influence of different TJ components on collective migration. While knockout of the transmembrane protein occludin has been shown to severely compromise migration dynamics,^[33]^ interference with the scaffolding ZO (zonula occludens) proteins was associated with both migration acceleration (ZO2; Raya-Sandino et al.^[34]^, ZO1; Bazellières et al.^[28]^) as well as deceleration (ZO1; Tornavaca et al.^[5]^, ZO3; Bazellières et al.^[28]^) in different epithelial cell lines. This discrepancy in evidence might be explained by the fact that such studies focused on the modification of only one single TJ component at a time. More recently, there were in-depth efforts to understand the impact of interfering with multiple ZO proteins on cell- and mechanobiology in general.^[1,2,4]^ However, the consequences for collective cell migration remain elusive.

To close this gap in knowledge, we performed migration experiments with ZO1 KO (single knockout) as well as ZO1/2 dKD (double knockdown) MDCK II cell lines as well as co-cultures comprising dKD and wildtype cells in a1:1 ratio accompanied by mechanical measurements and various imaging techniques. We showed that loss of ZO proteins substantially diminishes migration speed and coherence. This was induced by a liquid-to-solid-like transition leading to cellular jamming upon progressing migration. We found that collective cell migration is impaired by jamming through a mechanical tug-of-war that occurs in response to the enhanced contractility of MDCK II cells in the absence of ZO-1 and ZO-2. In the adherent state the dKD cells try to keep the balance between maximization of area occupied by the cells (adhesion) and assuming a highly contractile state induced by the enhanced perijunctional actomyosin ring. This leads to the coexistence of two subpopulations reflecting the interplay between cell-matrix and cell-cell adhesion. One subpopulation (‘winner’ cells) consists of small contractile cells that exert enormous stretching forces on their neighboring cells. As a result, a second population (‘loser’ cells) emerges displaying a larger, elongated footprint and increased apical tension. The increase in tension fosters cell division preferentially among the large ‘loser’ cells, which in turn feeds the subpopulation of small condensed ‘winner’ cells. In the final analysis, this fosters cellular jamming and thus renders the entire monolayer less mobile. Stalling cellular division in the later stage of confluency reestablishes faster migration since jamming is scaled down.

ZO1 KO cells showed similar but less pronounced proliferational, mechanical, and cytoskeletal adaptations. Albeit they also exhibited signs of jamming at late migration stages, no distinct small and large cell phenotypes arose as found for dKD cells. This finding emphasizes that the coexistence of large cells, which proliferate more and induce crowding, and small cells, which migrate less actively, is an important feature of jamming in TJ-deficient cells.

## 2. Results

### 2.1. ZO proteins ensure fast and coherent epithelial migration

To investigate the role of TJs in collective cell migration, we first performed migration experiments using phase contrast microscopy combined with particle image velocimetry (PIV)-based analyses (**Figure 1A/B**).^[35]^ Strikingly, video microscopy revealed that migration velocity of dKD cells was substantially lower than that of WT (wildtype) cells and even ZO1 KO cells. We first summarized data from the overall migration dynamics of the whole cell layers by averaging over all time points and all vectors (Figure 1C). While ZO1 KO cells did not display significant changes in migration dynamics (16 ± 2 µm h^−1^ (mean ± s.d.)) compared with WT MDCK II (18 ± 2 µm h^−1^ (mean ± s.d.), *p* = 0.13), ZO1/2 dKD cells migrated significantly slower (8 ± 1 µm h^−1^ (mean ± s.d.), *p* < 0.001). Additionally, we calculated the order parameter, which quantifies how directed the local motion is towards the migration edge (Figure 1C). We found that dKD cells migrated less directed (order parameter of 0.12 ± 0.04) than WT (0.29 ± 0.09) and ZO1 KO (0.31 ± 0.08) cells (*p* < 0.001), respectively.

**Figure 1.**
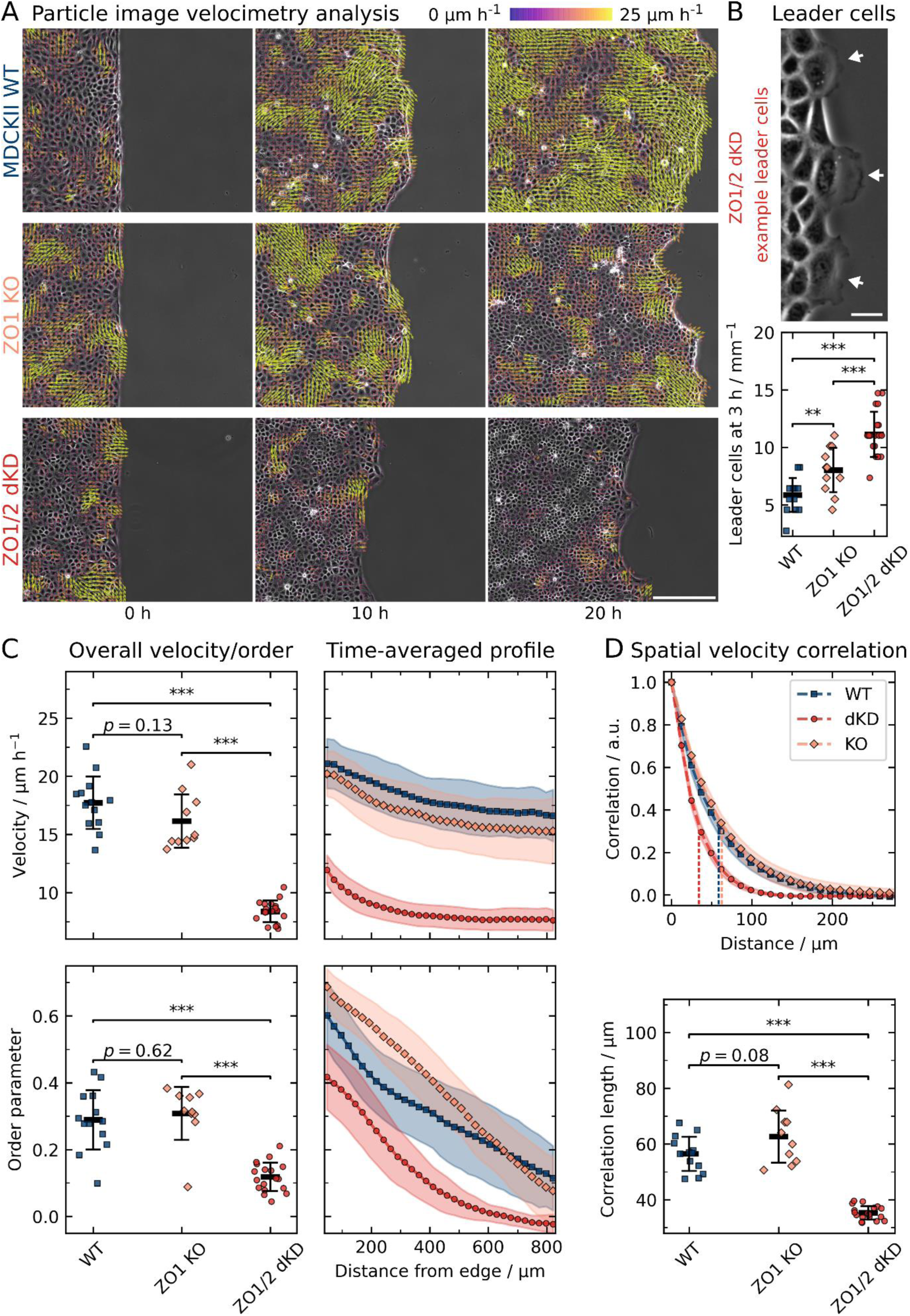
Collective cell migration dynamics of wild type (WT), ZO1 knockout (KO) and ZO1/2 double knockdown (dKD) MDCK II cells. A) Migrating cell monolayers with the corresponding velocity vectors obtained from particle image velocimetry (PIV). To enhance the figure’s visibility, cropped images are shown (about a fourth of the original field of view). Scale bar: 200 µm. B) Quantification of leader cell emergence and a corresponding dKD example. The amount of leader cells was normalized by the respective migration edge length for better comparison. Scale bar: 25 µm. C) The overall velocity and order are defined as the average over all vectors and time points, velocity and order were additionally averaged over time along the distance from the edge of the cell layer. D) Spatial velocity function. Vertical dashed lines indicate the corresponding characteristic correlation lengths below. All data are shown as means and standard deviations. Sample sizes (independent experiments): 13 (WT), 10 (KO), 18 (dKD).

To characterize the velocity transmission from the migration edge into the bulk of the monolayer, time-averaged velocity and order profiles were computed (Figure 1C). Here, we observed a subtle velocity decay in the range of the standard deviation with increasing distance from the migration edge for the WT and the ZO1 KO cells from about 20 µm h^−1^ at the edge to 17 µm h^−1^ (15% decrease) 400 µm away from the edge, while the dKD cells showed a sharper velocity drop from approximately 12 µm h^−1^ to 8 µm h^−1^ (33% decrease), approaching a plateau at about 400 µm, indicating an impaired velocity transmission from the edge into the layer. The order parameter decreased with increasing distance from the edge into the bulk layer for all three cell lines. Interestingly, for dKD cells the order parameter was not only lower at every distance from the edge but even approached zero at approximately 600 µm (indicating zero net movement towards the edge). This highlights that the cell collectivity was diminished, which goes hand in hand with the increased number of leader cells emerging from the dKD layers (Figure 1B). Almost twice as many leader cells were observed in the dKD (11 ± 2 mm^−1^ (mean ± s.d.)) as in the WT monolayers (6 ± 2 mm^−1^ (mean ± s.d.), *p* < 0.001). The ZO1 KO cells also showed an elevated number of leader cells (8 ± 2 mm^−1^ (mean ± s.d.), *p* < 0.01) compared with the WT, albeit less leader cells than the dKD variant.

To further study the reach of force transmission into the monolayer, we also computed the spatial velocity correlation of the migrating cells (Figure 1D). While the spatial velocity correlation of ZO1 KO cells decayed slightly slower than that of WT cells, yielding longer correlations lengths of 63 ± 9 µm (mean ± s.d.) for the KO than 57 ± 6 µm (mean ± s.d.) for the WT (*p* = 0.08), dKD cells showed considerably shorter correlation lengths of 35 ± 2 µm (mean ± s.d.) than both WT and ZO1 KO cells (*p* < 0.001, respectively).

Taken together, these findings suggest that ZO1/2 dKD cells migrate slower, less correlated, and less directed than the WT, thereby showing a significant loss of the hallmark parameters of cell collectivity. This behavior could be induced by a variety of mechanisms, from biochemical signaling to cell mechanical adaptations and possibly cellular jamming, in which the last two ones will be investigated further (*vide infra*).

### 2.2 ZO proteins prevent jamming and cell crowding

The following question remains to be answered: To what extent is the reduced migration speed an intrinsic property of ZO1/2-depleted cells or a consequence of compromised cell-cell contacts and therefore a collective effect? **Figure 2**A shows the analysis of both single cell migration and migration of individual cells within a confluent monolayer. Interestingly, the velocity of single dKD cells in the absence of cell-cell-contacts is even larger than the velocity of single WT cells (21.5 µm h^−1^ compared with 15.3 µm h^−1^ (median, *p* < 0.01)), which is in sharp contrast to their collective behavior when being part of a confluent monolayer. To scrutinize this collective effect, we expanded our time-averaged PIV analysis from before (chapter 2.1) to cell tracking in monolayers during very early (first 0.5-2.5 h) and late (19-21 h) migration. At an early confluent stage, the difference between wildtype cells and dKD is moderate (16.3 µm h^−1^ compared with 11.0 µm h^−1^, *p* < 0.001), while the drop in migration speed becomes more pronounced at a later stage when jamming sets in (*vide infra*). We rationalize the increased single-cell motility of dKD cells by their increased contractility and actomyosin activity due to upregulation of ROCK.^[36,37]^ We can therefore safely rule out that dKD cells are intrinsically less motile, rather the opposite. In the early stage (Figure 2A), where proliferation is largely absent, likely the enhanced contractility of dKD cells as described below and by previous studies already slows down collective migration compared to WT cells.^[1,2,4]^ Already from visual inspection of the epithelia, it was obvious that the KO and particularly the dKD monolayers showed decreased contact inhibition, becoming increasingly dense over time during migration due to continuous proliferation, whereas the WT layers showed no obvious change in density. Therefore, we quantified this peculiarity and also examined the impact of crowding on collective migration. While PIV is a well-established technique for the quantification of migration dynamics of cell collectives, it lacks information about the behavior of individual cells in the layer. To overcome this limitation, we applied the automated cell segmentation algorithm Cellpose (Stringer et al.) outlining the area occupied by each individual cell in 2D as shown in Figure 2B.^[38]^

**Figure 2.**
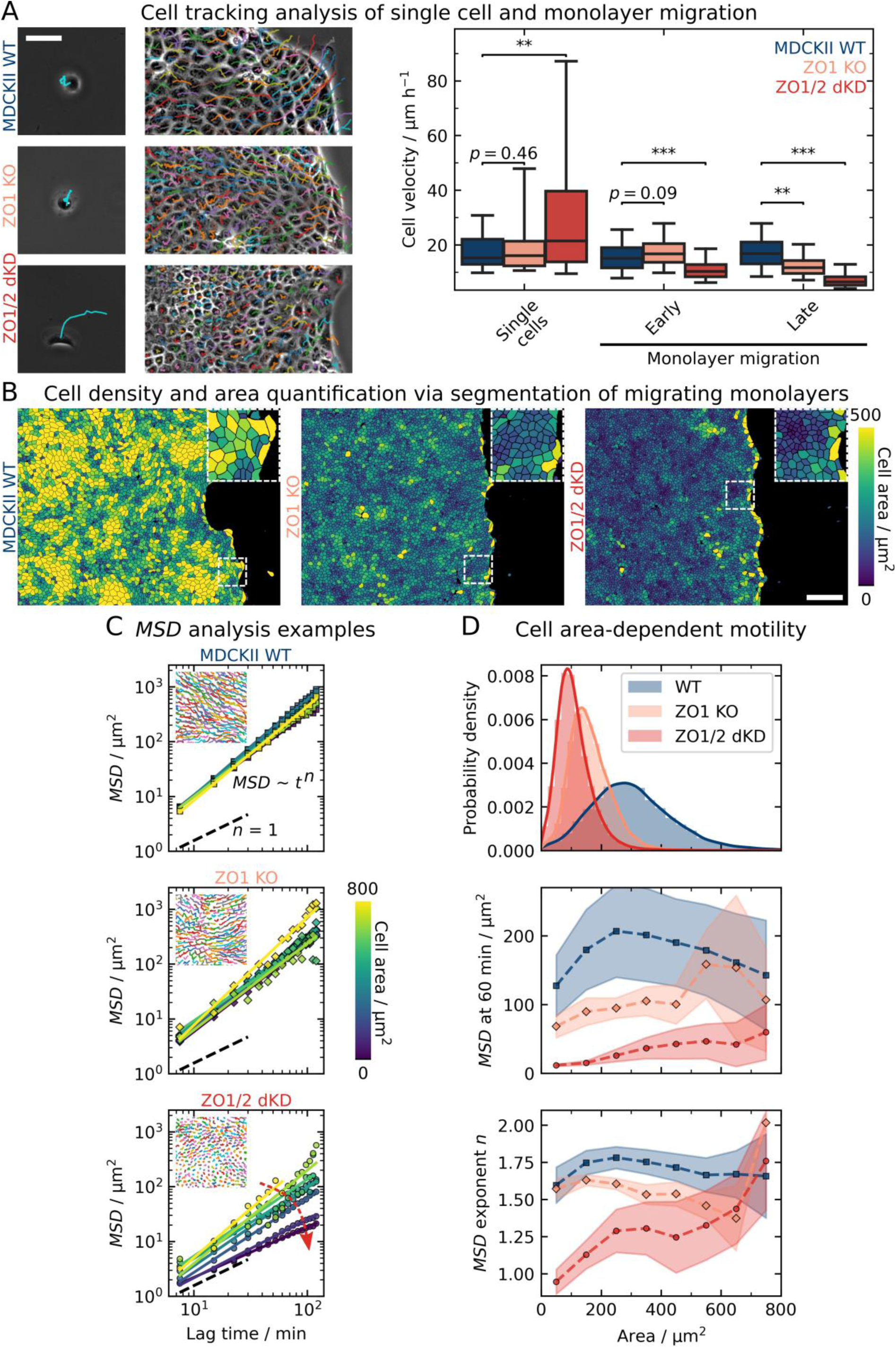
Quantifying individual cell velocities of early and monolayer as well as single cell migration using tracking and segmentation-based cell area-dependent motility analysis. A) Migration tracks of single cells and individual cells in a confluent cell monolayer colored randomly and quantification of the average velocity of individual cells during single cell migration and during early (0.5-2.5 h) and late (19-21 h) monolayer migration. Scale bar: 50 µm. B) Segmented cells in migrating monolayers after 20 h of migration, colored by the respective individual projected area in 2D. Scale bar: 200 µm. C) Tracking and mean-square-displacement (*MSD*) analysis. *MSD*s and corresponding power law regressions for a time window between 19 h and 21 h are shown for ensembles of cells in 100 µm^2^ area bins of an exemplary movie per cell line. The insets show exemplary tracks colored randomly. The red arrow indicates a decrease of the *MSD* with decreasing area, only prominent for the dKD cells. D) Cell area-dependent *MSD* parameters from C. The area distribution of all cells after 20 h of migration is shown at the top. ZO protein interference induced a shift to smaller areas with a pronounced skewness of 1.23 for the KO and 2.25 for the dKD as compared with the WT cells (0.94). Below, the *MSD* at 60 min and the power law exponent *n* are plotted vs the cell area. Points correspond to bins of 100 µm^2^ (around the point location), starting from 0 µm^2^. Sample sizes: 13 WT, 9 KO, 18 dKD independent monolayers, and 53 WT, 54 KO, 141 dKD cells (distributed over 2 independent experiments each). The boxes in A show the median and upper and lower quartiles. Whiskers indicate the 5^th^ and 95^th^ percentile. The boxplot comprises individual cell velocities. For more accurate statistical testing, cell velocities were averaged over separate monolayers. Means and standard deviations are shown in D.

Indeed, we found a substantial lack of contact inhibition of proliferation for the dKD cells as indicated by a strong increase of the cell density over time during migration (Figure S1A). While the density of the WT cells remained approximately constant, the KO cells displayed a cellular density increase similar to dKD cells but less pronounced. Yet, this density increase could also come from a lack of edge displacement combined with additional cells moving into the field of view. To confirm that mainly proliferation induced the density increase, we quantified cell density without a migration edge in a separate experiment (Figure S1E). Indeed, the dKD cell density increased stronger within the first 60 h and then reached a higher steady-state density than either WT or KO cells.

Two prominent parameters serve to characterize jamming transitions of cell layers: cell density and cell shape. Particle-based models attribute jamming to an increased cell density,^[39]^ whereas vertex models predict the shape of cells, as quantified by the shape index or the projected aspect ratio in 2D, to be the main determinant for jamming.^[24]^ However, along with the density increase with elapsed time, we did not observe a clear change in the projected cell aspect ratio in 2D (length divided by width) as shown in Figure S1A. Except for a short increase to a median aspect ratio of 1.60 around 5 h for the WT, all cell lines had a similar and only very subtly decreasing aspect ratio at around 1.45. However, the WT cells exhibited a slightly higher aspect ratio at all times, with a slightly broader distribution shifted to larger values (Figure S1B). Note that the observed aspect ratio values here are above the jamming threshold of 1.18,^[21]^ as calculated from the shape index of 3.81 as previously proposed by Bi et al.^[24]^ Notably, there was no correlated variation between cell area and aspect ratio of individual cells (Figure S1C), rendering these parameters largely independent of each other for each cell.

The decrease of migration velocity over time of dKD cells together with their increased proliferation rate suggests that jamming of the monolayer slows down migration speed. It is, however, important to distinguish earlier stages, in which cell division is still absent, from later stages, in which jamming increases due to increased proliferation. In the early stage, less mobile clusters of small contractile cells coexist with larger, highly strained cells in response to a competition between contractility and extensibility of cells to cover the matrix. The small contractile cells barely move and thereby slow down the monolayer. In later stages, the larger cells generate excess cells through cell division and thereby trigger jamming (*vide infra*).

A marked difference in the averaged PIV data of WT and ZO1-KO became apparent only after 15 h (Figure S1A), when the KO also slowed down and showed uncontrolled proliferation and slightly decreasing aspect ratios, similar to the behavior of dKD cells. However, the dKD cells display the slowest dynamics of all three cell lines, which could not be attributed solely to a change in the cell density, as this was also altered in ZO1-KO cells. The elevated contractility of dKD cells, which emerges in response to lack of ZO1/2 proteins, fostering remodeling and strengthening of the perijunctional actomyosin belt, is the key difference (*vide infra*).

So far, our purely mesoscopic and cell-averaged approach reveals a morphological heterogeneity (small and large cells) particularly for dKD cells at later time points (Figure 2B). This brought up the question, whether these morphological differences could be responsible for the impaired cellular dynamics. Therefore, we utilized single cell tracking to investigate the dynamics of individual cells in a layer during late-stage migration (19-21 h after insert removal), depending on the cell density and the projected cell area in 2D. We found that the motility, as quantified by the *MSD* (mean square displacement), of WT and KO cells generally did not depend on the cell area. In contrast, the *MSDs* of individual dKD cells show a clear dependency on cell area (Figure 2C). Specifically, we observed that the movement amplitude (*MSD* at 60 min) as well as the exponent *n* of the *MSD*s as a function of lag time rises with increasing cell area for the dKD. The small and most abundant bulk cells with an area around the distribution peak of about 120 µm^2^ showed passive diffusion-like movement with *n* ∼ 1 and small amplitudes of about 10 µm^2^. In contrast, the larger cells exhibited active motion with up to *n* = 1.75, which is similar to the WT cells and close to straight-line motion at *n* = 1.75, and five-fold increased amplitudes of 50 µm^2^ (Figure 2D). The KO cells had a similarly skewed cell area distribution with small bulk cell of about 180 µm^2^ showing movement amplitudes of about 80 µm^2^ while the sparse large cells moved about 110-200 µm^2^. However, neither WT nor KO cells showed any clear dependence of *n* on the cell area. Interestingly, the WT showed a more symmetrical cell area distribution around 280 µm^2^ (skewness of 0.94 as compared with 1.23 for the KO and 2.25 for the dKD cells) with averaged-sized cells showing the largest movements (*MSD* around 200 µm^2^) and cells at the extreme ends of the distribution moving less (*MSD* of about 130 µm^2^). Importantly, we did not find any clear dependence of the individual cell motility on the aspect ratio (Figure S1B).

Together, these results show that contractility and cell density, the latter being the result of the former, are the determining factors explaining the observed jamming of dKD cells due to an abundance of slow-moving small cells coexisting with faster-moving large and actively dividing cells.

As we have identified an important connection between jamming, proliferation, and migration speed in cells lacking ZO proteins, we investigated the distribution of the Yes-associated protein (YAP), a Hippo mechanotransduction signaling effector localizing to both the cytoplasm and the nucleus, where it is in its active state. YAP is known to influence cell proliferation as an important effector of the Hippo pathway, playing important roles in regulating cell migration. Mechanical signals that regulate YAP nuclear import comprise pulling, compressing, and shearing detected through cell-substrate as well as cell-cell junctions but also cytoskeletal remodeling.^[40,41]^

Therefore, we determined the ratio of YAP found in the nucleus to YAP in the cytoplasm for wildtype and dKD MDCK II cells (**Figure 3A**). We found indeed that in dKD cells a higher relative amount of YAP (0.8 ± 0.2 (median ± s.d.)) transits into the nucleus than in WT cells (0.5 ± 0.2 (median ± s.d.), *p* < 0.001), suggesting an increased propensity for proliferation in ZO1/2 depleted cells. As discussed above and elaborated further in the following, uncontrolled contractility eventually leads to jamming and consequently slows down collective cell migration. Additionally, the interplay of contractility and proliferation generates jamming through a positive feedback loop in which the small contractile cells stretch their neighbors to trigger cell division that in turn increases the number density of small, contractile cells (see below and Figure S7). Eventually, the monolayer assumes a frustrated state and slows down like a glassy material.

**Figure 3.**
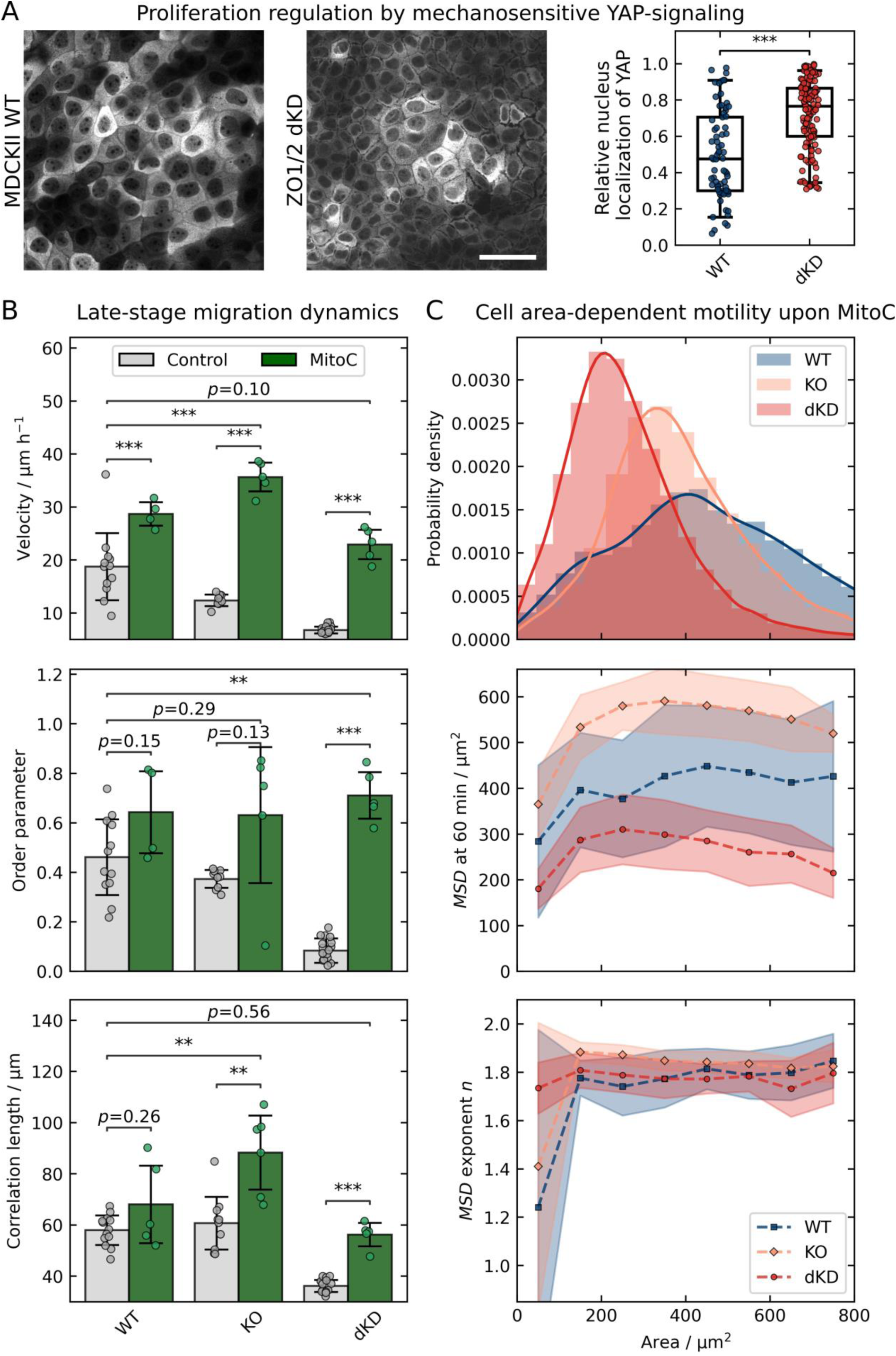
Late-stage jamming is induced via Yes-associated protein (YAP)-based upregulation of proliferation and can be largely prevented by proliferation inhibition with Mitomycin C (MitoC). A) Confocal images and corresponding relative localization of YAP in the cytoplasm and nucleus of WT and dKD cells, respectively. Scale bar: 50 µm. B) Migration dynamics after 19 h with (control) and without proliferation (MitoC). Overall velocity, order and correlation were calculated as in Figure 1. C) Cell area-dependent *MSD* parameters upon proliferation inhibition (MitoC treatment). The area distribution of all MitoC-treated cells at 20 h of migration is shown at the top. Larger cell areas and skewness parameters of 1.01 (WT), 1.36 (KO), and 1.83 (dKD) were observed. Below, the *MSD* at 60 min and the power law exponent *n* are plotted vs the cell area. *MSD*s and corresponding power law fits were calculated for a time window between 19 h and 21 h, in accordance with Figure 2. Points correspond to bins of 100 µm^2^ (around the point location), starting from 0 µm^2^. Boxes in A show the median and upper and lower quartiles. Whiskers indicate the 5^th^ and 95^th^ percentile. Data in A correspond to 72 WT and 124 dKD cells. Means and standard deviations are shown in B and C.

We therefore aimed to stall or even reverse the cellular crowding and jamming by the inhibition of proliferation using the well-established drug Mitomycin C.^[20,22,42–44]^. As expected, upon Mitomycin C treatment, the density of all three cell lines did not increase but instead even slightly decreased over time, confirming a successful inhibition of proliferation (Figure S2A). Concomitantly, the migration velocity increased while the overall aspect ratio slightly decreased over time. To quantify the impact of proliferation inhibition, we now focused again on the late-stage migration dynamics after 19 h (Figure 3B).

The drug increased the migration speed of WT MDCK II cells from 19 ± 6 µm h^−1^ to 29 ± 2 µm h^−1^ (mean ± s.d., *p* < 0.001), whereas the order parameter and correlation length did not change significantly (*p* = 0.15 and *p* = 0.26, respectively).

In comparison, we observed a significant increase of all migration parameters for the dKD cells (Figure 3B). Specifically, the dKD velocity increased from 7 ± 1 µm h^−1^ to 23 ± 3 µm h^−1^ (mean ± s.d., *p* < 0.001), which is similar to the velocity of untreated WT cells (*p* = 0.10), the order increased from 0.08 ± 0.05 to 0.7 ± 0.1 (mean ± s.d., *p* < 0.001), which is significantly higher than that of untreated WT cells (0.5 ± 0.2 (mean ± s.d., *p* < 0.01)), and the correlation length increased from 36 ± 2 µm to 56 ± 5 µm (mean ± s.d., *p* < 0.001), which is similar to the correlation length of untreated WT cells, being 58 ± 6 µm (mean ± s.d., *p* = 0.56).

The ZO 1 KO cells showed a similar behavior as the dKD cells upon proliferation inhibition, but with a less pronounced increase in all parameters. The velocity of KO cells increased in the presence of Mitomycin C from 12 ± 1 µm h^−1^ to 35 ± 3 µm h^−1^ (mean ± s.d., *p* < 0.001), which is also significantly higher than the velocity of untreated WT cells (*p* < 0.001), the order parameter increased from 0.37 ± 0.04 to 0.6 ± 0.3 (mean ± s.d., *p* = 0.13), which is slightly higher than the order parameter of untreated WT cells (*p* = 0.29), and the correlation length increased from 61 ± 10 µm to 88 ± 14 µm (mean ± s.d., *p* < 0.01), which is also higher than the correlation length of untreated WT cells (*p* < 0.01).

Taken together, the velocimetry data showed that inhibition of proliferation largely prevented the very late jamming process of dKD, and, less pronounced, that of ZO1 KO cells, by preventing an uncontrolled density increase.

Interestingly, the area-dependence of the *MSD* of dKD cells during late migration (19-21 h, *vide supra*) also vanished upon inhibition of proliferation (Figure 3C). In general, the individual cell area was larger in the presence of Mitomycin C for all three cell lines as expected for proliferation inhibition. This was most pronounced for dKD cells, where the cell area increased from 120 µm^2^ to about 220 µm^2^ (see area distribution in Figure 3C). Importantly, less separation into small and large cells occurred as indicated by the decreased area distribution skewness (Figure 3C) and confocal side views (Figure S2C). The *MSD* also showed a higher amplitude (*MSD* at 60 min) as well as exponent *n* for all treated cell lines than for untreated cells. Specifically, the dKD *MSD* amplitude was between 200 µm^2^ and 300 µm^2^ upon proliferation inhibition, which is slightly higher than for the untreated WT cells (150-200 µm^2^). The treated WT cells showed slightly larger amplitudes of 300-400 µm^2^ and the KO cells surpassed both other cell lines at about 300-550 µm^2^. Interestingly, the *MSD* exponent is equalized upon proliferation inhibition for all cell lines at about 1.8, which indicates completely restored directionality. This is in good accordance with the higher order parameter upon proliferation inhibition as shown in Figure 3B (*vide supra*).

Notably, upon proliferation inhibition we did not observe a clear trend in the movement amplitude and the *MSD* exponent with decreasing cell area as before anymore (Figure 3C). In contrast to untreated KO and dKD cells, upon inhibited proliferation the KO and dKD bulk cells of about 380 µm^2^ and 220 µm^2^, respectively, even showed a peak in the movement amplitude, while larger and smaller cells both moved slightly less. The *MSD* exponent remained constant with varying area.

Together, these results indicate that the strong crowding-induced jamming during late migration could be prevented by proliferation inhibition. However, it is important to note that, besides proliferation, Mitomycin C might also influence other cellular functions, which could potentially contribute to the observed migration dynamics. It is conceivable that since the propensity of large cells to divide is abolished, less excess volume is generated by the ‘loser’ cells and thereby the pulling of the small cells stalled. This leads to cells almost equal in height as shown in Figure S2C.

### 2.3 Successful ZO knockdown induces actomyosin remodeling

Given such severe phenotypical changes in the migration dynamics and proliferation rates of the ZO1 KO and dKD cells, we sought to investigate the phenomena also on the molecular level. First, to ensure successful genetic knockout, we performed confocal immunofluorescence microscopy. Indeed, ZO1 and ZO2 proteins were no longer visible upon double knockdown (Figure S3). Corresponding western blot analyses can be found in Beutel et al.^[45]^

ZO1 knockout was also successful as shown in Figure S3A. Importantly, ZO2 was only slightly upregulated indicating a possible compensation for ZO1. Notably, adherens junctions are not obviously affected (Figure S6) highlighting that the observations described here mainly reflect the ZO protein loss.

Since the transmembrane proteins in tight junctions are connected to the actin cytoskeleton via ZO proteins, we next investigated changes in the actomyosin architecture of the cells (**Figure 4**). Indeed, the actin cytoskeleton of the dKD cells was changed in a distinct way as shown in Figure 4C. Actin was accumulated at the periphery of individual cells, organized in thick rings, which were slightly separated at the apical plane of neighboring cells. ZO1 knockout cells on the other hand showed an intermediate phenotype with a less marked actin accumulation at cell-cell borders with a slight separation into two thinner cables. In comparison, the WT cells displayed the typical actin structure of MDCK II cells with a continuous mesh between cells and without any separation between neighboring cells or any obvious actin accumulation.

**Figure 4.**
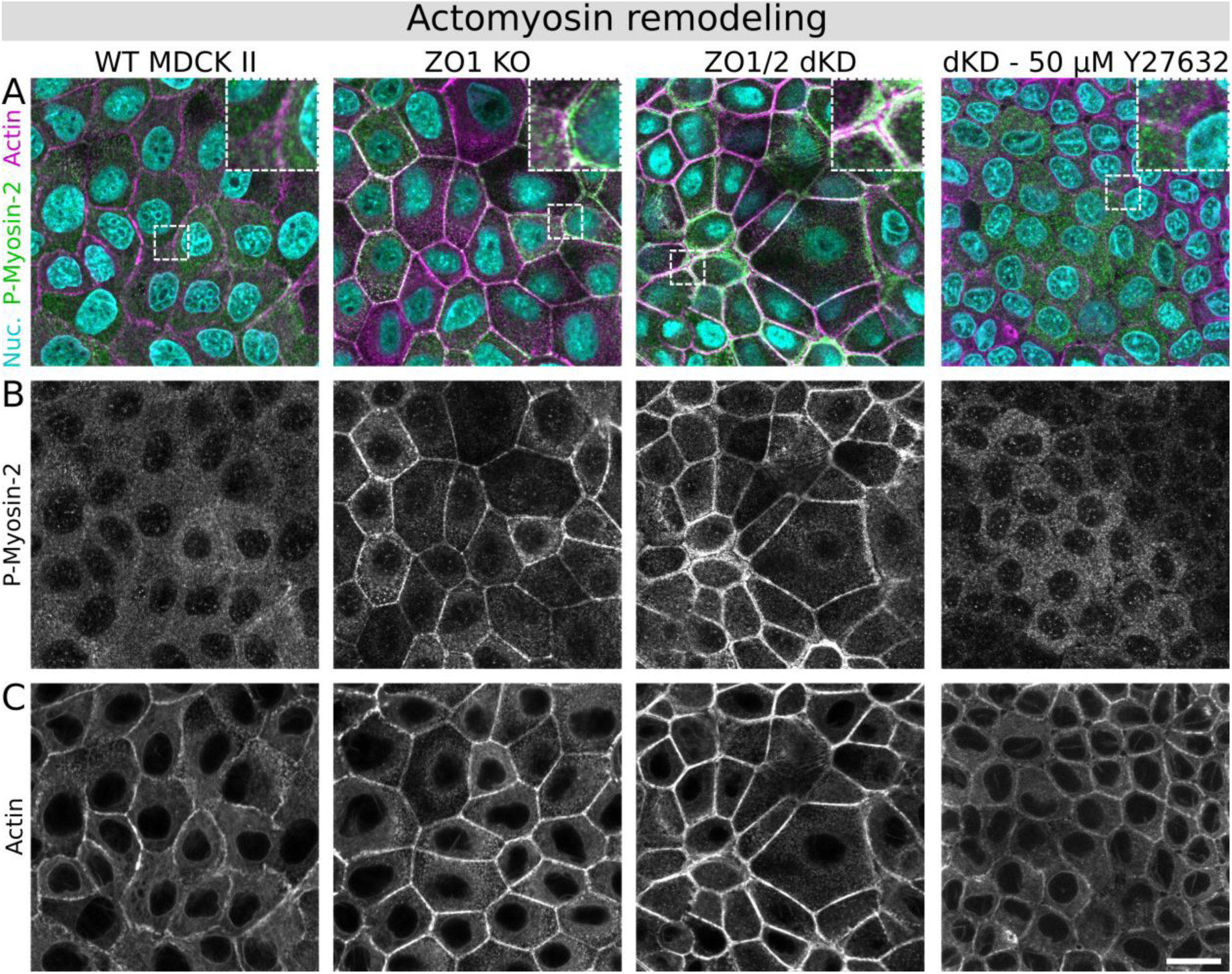
Actomyosin architecture remodeling upon ZO protein interference. A) Phosphorylated Myosin-2 (P-Myosin-2; green), actin (magenta) and nuclei co-staining of all three MDCK II cell lines and 50 µM Y27632 treated dKD cells (22.5 h incubation upon late migration). B) Corresponding gray-scale images of P-Myosin-2. C) Corresponding gray-scale images of actin. Shown representative examples far from the edge of migrating monolayers. Scale bar: 20 µm.

In addition, activated (phospho-) myosin-2 upregulation was particularly prominent at the cell-cell border in conjunction with the actin accumulation in dKD cells (Figure 4B), indicating upregulated actomyosin contractility. Interestingly, it seems that smaller dKD cells accumulated more peripheral actomyosin than their larger neighbors. On the other hand, ZO1 KO also showed accumulation of activated myosin at the cell periphery, albeit not as severe as in the dKD. In contrast, the WT cells showed little activated myosin without any prominent pattern or structure. Additionally, the occurrence of many small and some large dKD cells (as described above) was observed. In contrast, the WT and KO cell area appeared much more homogeneous.

If contractility is the key feature for the observed emergence of two subpopulations of cells balancing their shape through a tug-of-war in keeping the balance between pulling forces and cell-matrix interactions we would expect to relax this highly tensed state when impairing actomyosin contractility by blocking Rho-ROCK signaling. Therefore, we used the ROCK inhibitor Y27632, a cell-permeable and highly selective inhibitor of Rho-associated, coiled-coil containing protein kinase (ROCK) to reinstall tension homeostasis in dKD cells. Figure 4 exemplarily shows that P-Myosin-2 upregulation and actin remodeling is reversed in dKD cells upon addition of the inhibitor. Additionally, the two subpopulations of dKD cells disappear upon actomyosin relaxation, supporting our hypothesis that mechanical imbalance is responsible for the emergence of two cell populations. Upon Y27632 administration, the cells clearly adopt a homogenous size in the monolayer, rendering almost indistinguishable from the morphology of the wildtype. Figure S5 provides further experiments also at lower Y27632 concentrations and an additional quenching experiments in which we supply Y27632 to a cell layer during later stage migration of a confluent monolayer.

Taken together, these findings show that interfering with ZO proteins induces actin remodeling accompanied by myosin activation and accumulation, which is reversible by ROCK inhibition. This suggests that lack of ZO proteins is directly responsible for formation of contractile cells entering eventually a jammed through a mechanical imbalance that generates condensed clusters of immobile cells.

### 2.4 The cell topography reflects actomyosin remodeling upon ZO knockdown and shows a heterogeneous apical cell height distribution

Because severe actomyosin remodeling and accumulation at the apical cell periphery was observed, we also expected changes in the cellular topography (**Figure 5**). Consistent with the changes in the actomyosin structures, using AFM (atomic force microscopy) imaging we found prominently elevated ring-like structures at the periphery of individual dKD cells, slightly separated from each other (zoom-in in Figure 5A). In contrast, wild type cells exhibit a less pronounced but continuous cell border. ZO1 KO cells showed only a slight change of the cell border topography.

**Figure 5.**
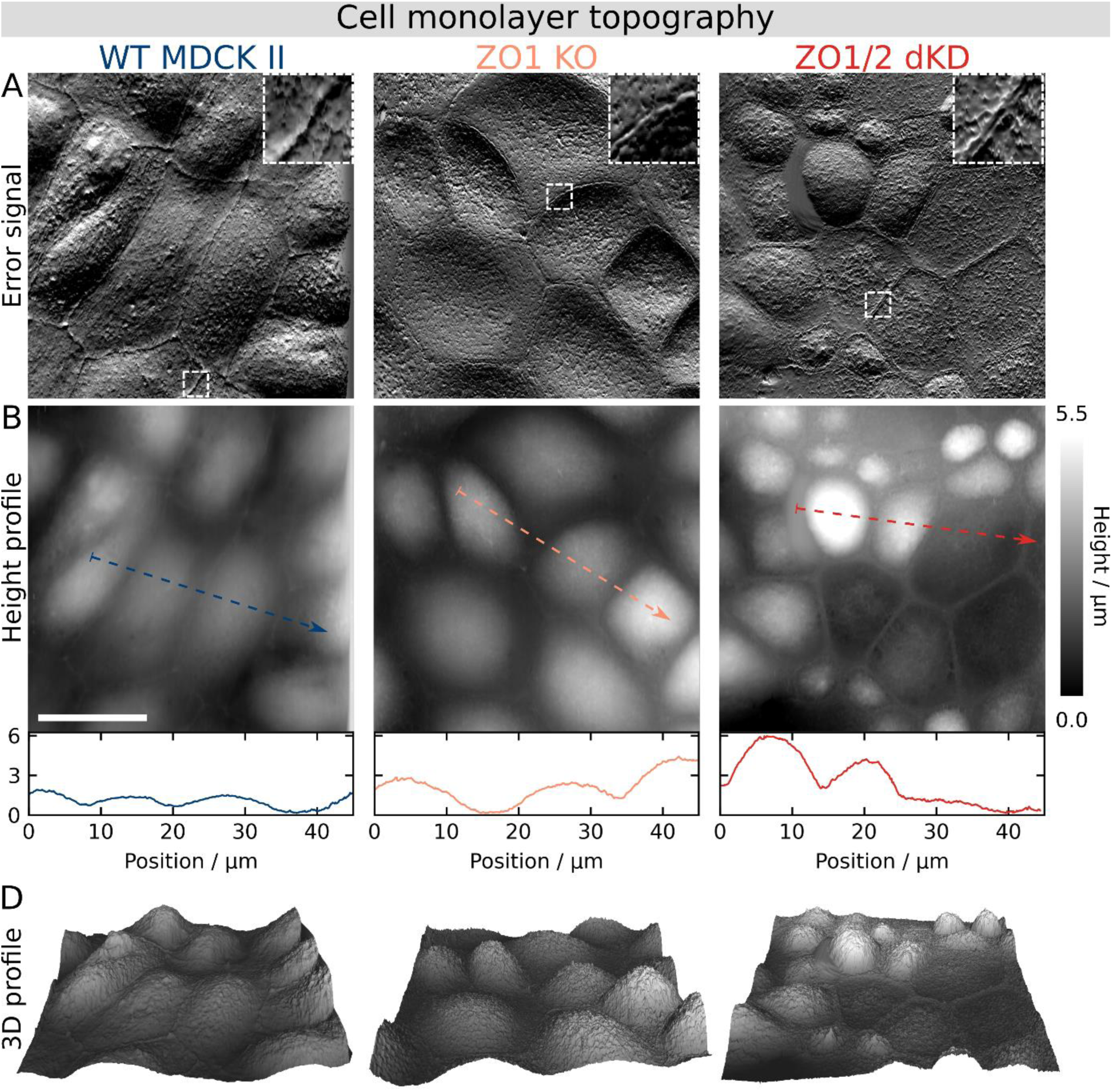
Cell monolayer topography adaptations reflect the actomyosin remodeling upon ZO protein interference as shown by AFM imaging. A) Error signal (deflection images). B) Height profile and cross-sections. D) Corresponding 3D topography maps, slightly up-scaled vertically, with the *z*-axis length being 20% of the *x*/*y*-axis (13.3% corresponds to an aspect ratio of 1). Scale bar: 20 µm.

Furthermore, AFM imaging confirmed the data from confocal fluorescence microscopy and segmentation indicating a pronounced height and area heterogeneity in dKD cells. While the apical cap of cells with a small area of about 100 µm^2^ (compare with chapter 2.2) was several micrometers high (> 3 µm), other cells were larger in area but do not exhibit any distinct apical cap rising above the peripheral ring. In comparison, the apical cap of WT cells was typically 1-1.5 µm high and homogenously distributed across the monolayer. The ZO1 KO cells displayed an intermediate phenotype with a homogenous cap height distribution, which are typically slightly higher than WT cells, at about 2-3 µm.

In conjunction with the actomyosin results, these data show that ZO1/2 dKD consistently induces distinct molecular and topographical changes, most notably, severe actomyosin accumulation underneath the membrane at the cell-cell borders in the small cell population being responsible for altered mechanical properties, which are scrutinized in the next chapter.

### 2.5 ZO proteins are necessary for mechanical integrity and tissue fluidity by preventing an uneven tug-of-war-like imbalance

In light of the prominent cell topography adaptations and concomitant actomyosin remodeling, and because contact inhibition of proliferation and jamming are typically tightly coupled with cellular mechanics, the consequences of ZO depletion for cell mechanics were investigated. To this end, we performed AFM measurements with an emphasis on force relaxation experiments that also permit to assess the rheological properties of the cells. First, force volume imaging showed that stiffness was increased considerably at the cell periphery of ZO1 KO and dKD compared with WT cells (AFM maps in **Figure 6A**), whereas the center appeared to be softer compared with WT cells.

**Figure 6.**
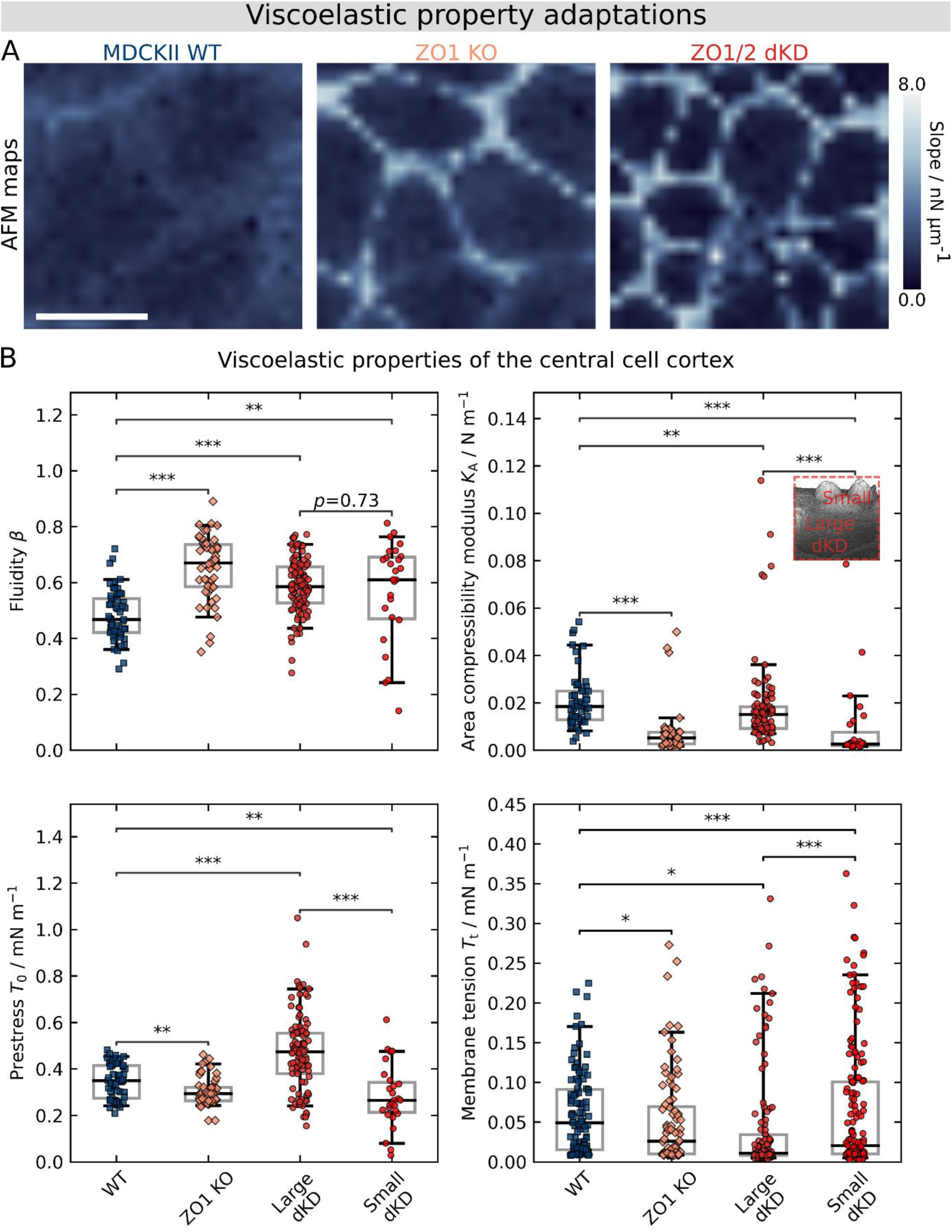
ZO proteins ensure viscoelastic integrity of cells as shown by AFM (atomic force microscopy). A) Exemplary AFM maps of migrating WT, ZO1 KO and ZO1/2 dKD cells showing the slope of the force during contact, mirroring the apparent stiffness of cells. Scale bar: 20 µm. B) Site-specific viscoelastic properties of the central cell cortex. Fluidity *β*, area compressibility modulus *K*_A_, prestress *T*_0_, and membrane tension *T*_t_ are shown. Five curves were immediately recorded on the same position at the center of one cell. Individual data points represent the average of the respective fitting parameter for an individual cell. The boxes show the median and the upper and lower quartiles. Whiskers indicate the 5^th^ and 95^th^ percentile.

This is consistent with the observed accumulation of actin into a contractile actomyosin ring and the altered topography at the cell periphery of ZO depleted cells. Apart from stiffness maps, we also used site-specific indentation experiments followed by force relaxation to study the mechanical and rheological cortex properties in greater detail. The model we applied was introduced recently by Cordes et al.^[46]^ Briefly, it considers stress relaxation of the cortex according to a power law providing us with a prestress corresponding to the isotropic cortical tension *T*_0_ plus membrane tension *T*_t_, the area compressibility modulus *K*_A_ of the cortex and the fluidity *β*, which classifies the flowing propensity of the network. A *β* value of 1 corresponds to a Newtonian fluid whereas a value of 0 describes a solid. Since we observed a prominent heterogeneity of the dKD cell morphology with a flat surface observed for large cells and a high apical cap seen for smaller cells, we considered the resulting geometrical differences in the model and distinguished between large (about 200 µm^2^) and small dKD cells (80 µm^2^).

Notably, we observed statistically significant changes in all mechanical parameters upon ZO protein loss (Figure 6B).

Cortex-dominated prestress *T*_0_ was significantly lower in the center of KO (0.30 ± 0.06 (median ± s.d.) mN m^−1^, *p* < 0.01) and small dKD cells (0.27 ± 0.13 (median ± s.d.) mN m^−1^, *p* < 0.01) than in the center of WT cells (0.35 ± 0.07 (median ± s.d.) mN m^−1^), indicating a downregulation of the actin cortex in both populations due to remodeling of the actin cytoskeleton. In contrast, the large dKD cells showed an increased prestress (0.47 ± 0.16 (median ± s.d.) mN m^−1^, *p* < 0.001 compared with WT as well as with small dKD cells). This goes hand in hand with a flatter morphology indicative of area expansion, leading to higher tension. A similar behavior was found recently by us, in which the elastic modules of confluent MDCK II cells increase with increasing projected apical cell area in a nonlinear fashion. Generally, the prestress *T*_0_ contains contributions from i) membrane tension that originates from adhesion of the plasma membrane to the underlying cytoskeleton, ii) area expansion of the apical shell and iii) active contraction by myosin II motors. To tell apart the contribution of the actin cortex from that of the plasma membrane-cytoskeleton attachment to the prestress *T*_0_ we additionally pulled out membrane tethers upon retraction to measure the membrane tension *T*_t_. We observed that *T*_t_ decreased upon ZO protein KO for all cell lines. It dropped from 0.05 ± 0.05 (median ± s.d.) mN m^−1^ (WT) to 0.03 ± 0.06 (median ± s.d.) mN m^−1^ (KO, *p* < 0.05), 0.01 ± 0.09 (median ± s.d.) mN m^−1^ (large dKD, *p* < 0.001), and to 0.02 ± 0.08 (median ± s.d.) mN m^−1^ (small dKD, *p* < 0.05). This shows that the prestress changes were only partly explainable by a decrease in membrane tension. However, the membrane tension of large dKD cells decreased, supporting the idea that prestress of the larger and flatter dKD cells stems from area expansion rather than a reinforced attachment of the cortex to the membrane.

Along with smaller prestress, we also observed a fluidization of the cortex represented by an increase of *β* from 0.5 ± 0.1 (median ± s.d.) to *β* = 0.7 ± 0.1 (median ± s.d., *p* < 0.001) for KO cells, and to *β* = 0.6 ± 0.2 (median ± s.d., *p* < 0.01) for the small dKD cells, respectively. Also, for large dKD cells an increase in fluidity was found (*β* = 0.6 ± 0.1 (median ± s.d.), *p* < 0.001). Recently we showed that fluidity and area compressibility modulus of the cortex are not necessarily independent parameters. Accordingly, the area compressibility modulus *K*_A_ decreased from 0.02 ± 0.01 (median ± s.d.) mN m^−1^ for WT to 0.005 ± 0.001 (median ± s.d.) mN m^−1^ for KO (*p* < 0.001) and to even 0.003 ± 0.002 (median ± s.d.) mN m^−1^ for small dKD cells (*p* < 0.001), respectively. For the large dKD cells, *K*_A_ fell by only 50% to 0.01 ± 0.01 (median ± s.d.) mN m^−1^ (*p* < 0.01) albeit the fluidity was rather high (*β* = 0.6 ± 0.1). Notably, the large dKD cells show a significantly higher *K*_A_ than the small dKD cells. This might indicate the presence of a prestressed cortex with less membrane reservoir to compensate for the external deformation. This view is backed up by the finding that the geometrical apical membrane of the large dKD cells is also larger than that of the small dKD cells despite the apical bulging (as inferred from geometrical considerations based on the topography measurements in Figure 5). Interestingly, the large and prestressed dKD cells were observed to proliferate over twice as much as the small dKD cells (Figure S7), indicating a possible connection between the mechanical phenotype of the large dKD cells and proliferation.

The drop in area compressibility modulus in the small dKD and KO cells could be either due to a higher cortical elasticity or a larger apical excess area, giving rise to apparent area compressibility modules. Considering the substantial morphological changes of the apical membrane/cortex in response to ZO1/2 knock down, such as bulging of the cortex and the reported occurrence of membrane reservoirs (small dKD cells), it is conceivable that both effects contribute to the observed softening.

Taken together, these findings show a mechanical integrity loss upon ZO protein knockout: Actomyosin is recruited from the cortex to the periphery of individual cells building up a stiff and contractile actomyosin ring while leaving the apical cortex weakened. On one hand, this leads to bulging of the central cell cortex, formation of excess area, and fluidization of the cortex in small dKD cells, making these cells into ‘winner’ cells in this cellular tug-of-war. On the other hand, large ‘loser’ dKD cells are prestressed by the contractile small cells and thereby seem to start proliferating upon mechanical activation of the Hippo pathway through YAP signaling (Figure 3) and possibly Piezo1-signaling. In fact, a statistical analysis by visual inspection (Figure S7) confirms this hypothesis and shows that larger cells proliferate more frequently, while smaller ones divide less. As a consequence, a larger amount of the small ‘winner’ dKD cells, which exhibit jamming, are generated by the uncontrolled proliferation and gradually impair collective migration more and more.

### 2.6 The tug-of-war outcome is unequivocally determined in co-cultures of dKD and WT cells as a phase separation into two mechanically distinct subpopulations

Our working hypothesis proposes is that two subpopulations emerge in ZO1/2 depleted cells due to enhanced contractility as a result of cytoskeletal remodeling of the actomyosin belt. A way to ultimately verify this hypothesis is accomplished by substituting the larger ‘loser’ cells in the dKD layer by wildtype cells with intrinsically lower contractility. If our assumption was correct, we would expect a separation into small, highly contractile cells, exclusively consisting of dKD cells, coexisting with outstretched wildtype cells, replacing the former ‘loser’ cells. This shifs the tug-of-war to more dKD cells assuming a compact morphology due to higher contractility. As a consequence, jamming is introduced into the co-culture by the over-contractile ‘all-winning’ dKD cells and migration speed diminishes accordingly.

**Figure 7** shows the results of a co-culture analysis of a 1:1 mixture of WT and dKD cells at early and late-stage migration, respectively. WT cells that synthesize an intrinsic fluorophore (myosin-2 marked with GFP, green) were chosen to readily identify them in the co-culture. Figure 7A clearly shows that, after 20 h of migration, indeed two subpopulations emerge that can be unequivocally assigned to dKD cells (phase contrast with false color, magenta) and wildtype cells (green). After 20 h of migration, cells were allowed to migrate for another 6 h before the ROCK inhibitor Y27632 was added and incubated for 5 h (31 h overall migration, the last 5 h with 50 µM Y27632). This caused a dramatic relaxation and expansion of the dKD cells (Figure 7A) reducing the appearance of the two populations dramatically. During non-inhibited mix migration, the dKD cells are extremely small compared to the wildtype cells (see histogram, Figure 7B) and immobile rendering the whole layer less motile (Figure 7C). Indeed, our working hypothesis that over-contractile small cells are immobile and thereby slow down the whole layer, while larger extensile cells are still rather mobile, was confirmed.

**Figure 7.**
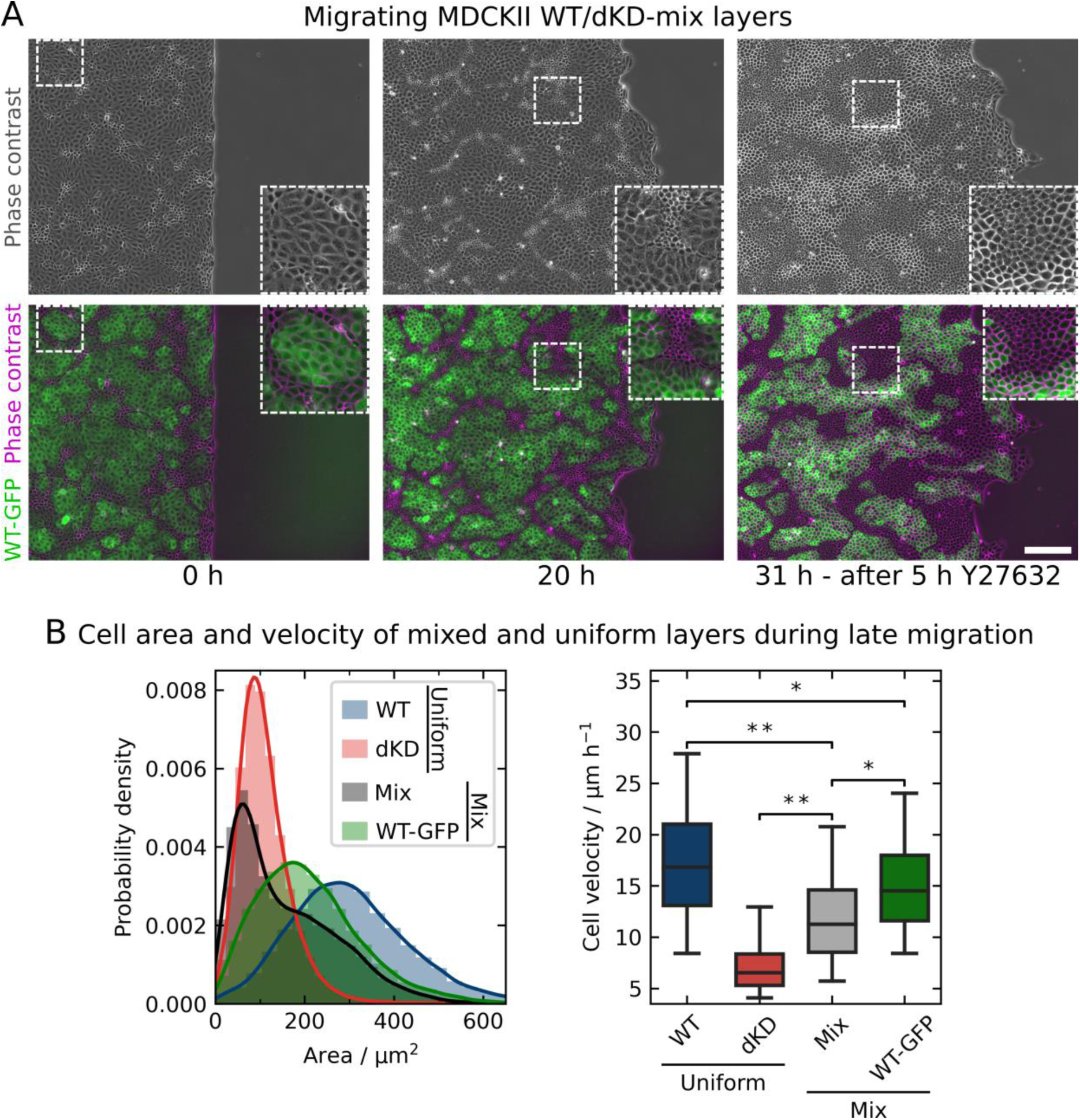
Results of co-cultured MDCK II WT and dKD cells mixed in a 1:1 ratio. A) Migrating cell monolayers examples of WT and dKD co-cultures at the start of migration, after 20 h of migration and after 11 h additional migration, the latest 5 h of which with 50 µM Y27632. WT cells (stably transfected with GFP-myosin-II-A, called WT-GFP) are shown in green, WT-GFP and dKD cells (phase contrast with false-color) are shown in magenta. Cropped areas shown for better visibility. Scale bar: 200 µm. B) Quantification of the cell area and velocities in pure WT and dKD and a co-culture of both cell types in monolayers during late-stage migration (area at 20 h and velocities between 19 h and 21 h). The left hand shows the area distribution of uniform WT and dKD cultures, as well as the 1:1 mix co-culture of both and the WT-GFP cells contained in the co-culture migration. Cell velocities on the right hand side of the same cell (sub-) populations have been calculated as in Figure 2. Samples sizes (independent monolayer): WT: 13, dKD: 18, Mix: 4. The histogram and boxplot comprise individual cells. For more accurate statistical testing, cell velocities were first averaged over separate monolayers.

Specifically, co-cultures moved significantly slower than pure WT cells (12.0 ± 1.2 µm h^−1^ compared with 18.3 ± 2.9 µm h^−1^ (median ± s.d.), *p* < 0.01), which was indeed slowed down by the contracted dKD cells. The generally higher motility of the wildtype cells maintained an overall larger migration velocity within the co-cultures compared to a cell layer solely consisting of dKD cells (12.0 ± 1.2 µm h^−1^ compared with 7.1 ± 0.6 µm h^−1^ (median ± s.d.), *p* < 0.01). However, WT cells in the co-culture were also slowed down due to the low-motility dKD cells (15.3 ± 0.8 µm h^−1^ (WT in mixed layer), 18.3 ± 2.9 µm h^−1^ (pure WT) (median ± s.d.), *p* < 0.05). Lastly, the WT cells in the co-culture mix were still faster than the overall mix (15.3 ± 0.8 µm h^−1^ compared with 12.0 ± 1.2 µm h^−1^ (median ± s.d.), *p* < 0.05). Accordingly, in the tug-of-war the ‘winners’ (dKD) slow down and induce jamming while the ‘loser’ (here WT) cells maintain a certain amount of fluidity and motility.

In summary, we found that addition of wildtype cells substituted the ‘loser’ cell population otherwise recruited from the dKD population, which is enforced in a tug-of-war between cell-cell and cell-matrix adhesion. This is distinct from phase separation based on differential adhesion between two cell types and has first been reported by Balasubramaniam et al. using wildtype MDCK II cells mixed with more contractile E-cadherin KO cells.^[47]^ The majority of dKD cells within this mixture is now capable of adopting the preferred, condensed and highly contractile phenotype, while the wildtype cells maintain their extensile behavior. The presence of this jammed phase eventually slows down collective migration of the whole layer and the wildtype. This series of experiments clearly supports our mechanistic view of the role of ZO-proteins not only as intracellular linkers that directly connect the actomyosin cytoskeleton with transmembrane adhesion proteins but also as regulators of apical tension. Once this delicate balance is perturbed by depletion of ZO proteins, a reorganization of the perijunctional actomyosin cytoskeleton into large thick cables of actomyosin occurs. Highly contractile and jammed cells emerge, which eventually lead to a partial jamming of the whole cell layer, exhibiting reduced migration speed.

## 3. Discussion

In this study, we were able to show that efficient collective cell migration depends on the tight junction ZO proteins. We show that ZO protein loss leads to severe cellular crowding and eventually jamming, which is fostered by morphological, mechanical, and cytoskeletal integrity loss.

Essentially, we found that ZO protein loss leads to formation of thick and contractile perijunctional actomyosin cables. This is in line with previous characterizations of cells lacking ZO proteins.^[1,4,8,11]^ Particularly, recent evidence suggests that TJs provide a negative mechanical feedback to the actomyosin cytoskeleton of individual cells in a layer, so that they do not contract and pull excessively.^[1]^ Because this feedback loop is missing in our cell lines, it is expected that most individual cells contract in an uncontrolled manner.

Indeed, many cells contract excessively via the perijunctional actomyosin ring. The constriction of this ring leads to laterally smaller cells with a projected area in 2D of about 80 µm^2^ that bulge out apically, presumably in order to maintain constant volume. Since actin is remodeled and potentially recruited from the cortex into these rings, the cortex is softened.

These observations are in line with recent studies showing similar actomyosin remodeling in conjunction with such morphological changes, particularly of the cell cap.^[1,4,8,11]^ In general, actomyosin remodeling is known to determine cell mechanical as well as morphological adaptations.^[48–50]^ Together, the dome-like apical membrane and the weakened cortex result in excess membrane area accompanied by lower prestress and higher fluidity while the actomyosin ring itself becomes extremely stiff as visible in our force maps.

In contrast to our observations of softening and fluidization of the cell body, former work by Cartagena-Rivera and coworkers report an overall tension and viscosity increase in ZO1/2 knockout cells.^[2]^ However, experiments in this study either targeted cell junctions directly or were carried out with much larger probes (> 20 µm) than our conical indenters of only a few tens of nanometers. Therefore, their measurements are integrated over a larger area capturing the mechanical response from both the extremely stiff cell borders and the soft cell body, which might explain the controversial findings.^[2]^ Another reason could be the fact that the authors used much longer cell growth times than us of over one week. Coupled with the uncontrolled proliferation, this might explain the discrepancies in the observed mechanical behavior: Upon long culturing times, the cell layer becomes increasingly dense and more small ‘winner’ cells, meaning more contractile cells and thereby actin rings per area which, in turn, will dominate the mechanical readout in those studies.

The balance between adhesion to the substrate or matrix and the intercellular tension leads to the coexistence of two subpopulations. Besides the small and contractile ‘winner’ cells a second population formed by large and outstretched ‘loser’ cells emerges displaying increased apical tension. This second cell phenotype occurs in both ZO1 and ZO1/2 depleted dKD cells. It is generally characterized by: 1) a larger projected area of 150-250 µm^2^, i.e., larger than most dKD cells but smaller than average WT cells, 2) thinner perijunctional actomyosin rings, 3) a flattening of the apical cortex, and 4) much higher prestress *T*_0_ and less excess membrane area than the dKD cells. Hence, two mechanically and morphologically distinct but coexisting dKD cell phenotypes emerge with time. For clarity, we distinguish between these two phenotypes and refer to them as *small (‘winner’)* and *large (‘loser’)* dKD cells.

The perijunctional actin contraction of the small cells is presumably responsible for the flattening and stretching of neighboring cells, which become larger. In response, the large cells need to sacrifice some of the excess area stored in the apical cell membrane, explaining the smaller decrease in *K*_A_ in contrast to the smaller dKD cells. Similarly, the pulling force from the contractile small cells is reflected in the increase in *T*_0_ in the large cells. Larger cells typically display larger tension due to lateral strain imposed from adjacent cells.^[51]^

In essence, the small cells contract and thereby pull on the large cells and stretch them balancing the forces across the cell layer.

However, the large cells are unable to escape from the tensile stress into the third dimension (which is further exemplified in 3D dKD cultures, where we do not observe a separation into small and large cells (data not shown)). As a consequence, the cells become laterally stressed and respond by proliferation (possibly by activation of the Hippo pathway through YAP signaling and/or though Piezo1)^[40,41,52]^, which relaxes the lateral stress. In turn, the increased proliferation leads to higher cell densities and eventually to partial jamming, impairing cell migration.

We could further confirm this view by virtually substituting the larger ‘loser’ cells with WT cells monitoring the dynamics of co-cultures comprising dKD and WT cells in a ratio of 1:1 (Figure 7). The co-culture with wildtype cells allowed all highly contractile dKD cells to adopt a condensed and contracted shape with an almost circular perimeter, while the WT cells took the place of the ‘loser’ cells being stretched and forming the liquid-like phase, which still displays higher motility. The dKD cells assume a jammed, solid-like phase, which slows collective cell migration of both cell populations substantially. This tug-of-war mechanism, where phase separation happens and ‘winner’ cells stretch out ‘loser’ cells is in line with quite similar co-culture experiments of contractile and non-contractile cells by Ladoux and co-workers.^[47]^ Essentially, we observe the same contractility-driven phase separation as a consequence of activity differences rather than differential adhesion between the two cell types. The observations of uncontrolled proliferation and mechanical imbalances are in line with the idea that TJs are both biological signaling hubs^[53,54]^ and mechanical sensors.^[1,3]^ Cell mechanics could be rescued by ROCK inhibition, identifying a crucial mechanosensitive pathway at play in ZO-depleted cells, which is in line with previous work.^[4,55]^ Notably, Matsuzawa et al. similarly observed distinct subpopulations in ZO1/2-depleted epithelial cells as well as WT/ZO-dKD mixes, which could be resolved by ROCK inhibition.^[55]^ Regarding proliferation, ZO proteins have been shown to directly control proliferation through cell cycle arrest.^[56,57]^ On the other hand, Rosenblatt and coworkers recently showed that mechanically stretched MDCK II cells divide more frequently than unstressed cells.^[52]^ Mechanical stretch itself rapidly stimulates cell proliferation through activation of the Piezo1 ion channel.^[52]^ We propose that the contractile smaller cells provide exactly this kind of mechanical stimulus leading to cell divisions primarily of the larger, flat cells, which are stretched considerably.

This proliferation and cell density increase coupled with the mechanical changes of individual cells leads to migration disruption and jamming. Strikingly, evidence accumulates that particularly smaller cells are responsible for the onset of jamming in dKD cells.

Specifically, the differences in the jamming expansion of KO and dKD cells, respectively, can only be explained by the area-dependence of active migration on the individual cell area. While the KO and WT cells display the same fast and active dynamics regardless of their cell area, dKD cells become increasingly less dynamic with decreasing cell areas. Particularly, the small dKD cells, which constitute the majority, show passive diffusion-like behavior with a power-law exponent of about 1, whereas the larger cells display active motion with a similar exponent to the WT cells of about 1.8. Accordingly, the small cells are particularly immobile and thereby impairing the migration of the whole dKD layer.

Together, these results draw the following picture (**Figure 8**): dKD cells respond to lack of tight junction proteins by remodeling of the perijunctional actomyosin into thick sarcomeric ring-like structure that contract excessively. The contraction leads to two coexisting subpopulations of cells: Small contractile cells pull on their neighbors, thereby generating so-called ‘loser cells’ that exhibit a large elongated footprint to maintain a maximized adhesion area. In response to external pulling they show an increased apical tension, since they are not able to escape into the third dimension (the two subpopulations are not observed in 3D cell culture). The large cells respond to the lateral stress by proliferation leading to even more small cells and eventually to crowding since a higher cell density allows the cells to transfer more and more cells into the subpopulation of small and immobile cells.

**Figure 8.**
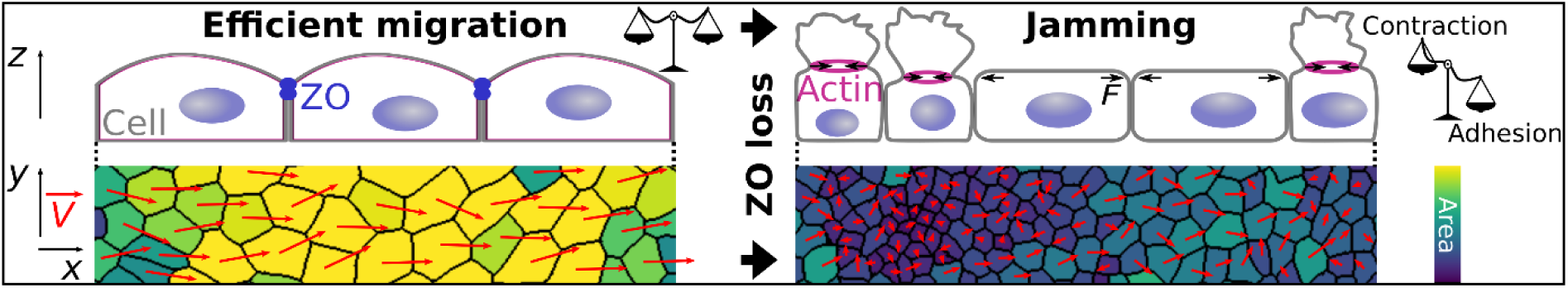
Proposed model of the delicate force balance necessary for cell layer fluidity. WT cells display a balance equilibrium between contraction and adhesion, and, thus, display a homogenous morphology and can migrate efficiently (left). In contrast, cells lacking ZO proteins develop a new and perturbed force balance leading to heterogeneous cell morphology and jamming (right). Two cell populations emerge: small, highly contractile cells with apically bulged-out excess membrane and large, stretched cells. The small cell population is particularly immobile and additional crowding amplifies this jamming.

On a mesoscopic scale, both migration velocity and order are diminished in ZO1/2 dKD cells and, upon progressing migration and proliferation, also in ZO1 KO cells. This is in line with recent work showing the deceleration of migration in cells lacking ZO proteins.^[5,58]^ For instance, endothelial cells lacking ZO1 were shown to migrate slower.^[5]^ For MDCK II cells, Fedele et al. found that the migration dynamics of monolayers with already inhibited adherens junctions are diminished upon ZO1/2 double knockout.^[58]^ In addition, ZO protein loss significantly shortens the spatial velocity correlation length. Along the same line, the KO and particularly the dKD monolayers develop more individual leader cells. Both velocity correlation and leader cell emergence were implicated as hallmarks of collective cell behavior and mechanical coupling.^[15,27,32,59,60]^ Accordingly, cells lacking ZO proteins behave less collectively and exhibit perturbed mechanical coupling.

Yet, since cell mechanics and proliferation are tightly coupled, the relative impact of each on the stalled migration remains to be elucidated. Therefore, we investigated the peculiar relationship between cell mechanics and cell density by inhibiting proliferation. In line with recent studies, proliferation inhibition slightly increases migration speed in WT MDCK cells.^[20,22]^ Strikingly, inhibiting the proliferation of dKD cells succeeds in almost complete recovery of the migration velocity, order, and correlation length observed for WT cells. This underlines the importance of the mechanically induced proliferation and cell crowding as a decisive control parameter for collective migration. However, it is conceivable that upon inhibited proliferation, the ‘loser’ cells cannot be stretched as easily anymore due to missing mitotic rounding forces, which might partly restore the mechanical balance.^[61]^

The observation that MDCK cells at low densities show such a high, and at higher densities a significantly lower power-law exponent is shared by recent experimental evidence.^[62,63]^ This general effect of cell density on collective migration dynamics is also in line with physical particle-based models of tissue dynamics.^[39]^ These models predict cell density to be the main determinant parameter for collective motility with motion arrest at high densities. However, to our knowledge, the direct dependence of cell motility of individual cells in a monolayer on their projected area has not been observed experimentally before.

Interestingly, the cell shape (projected aspect ratio in 2D), as predicted by vertex-based models, does not seem to be the decisive parameter in contrast to the cell area itself. Particularly, while we do see a slight shift towards lower overall aspect ratios, we do not observe a clear dependency of the motility of individual cells on the aspect ratio as on the area. It is important to note that instead of addressing the aspect ratio of individual cells, current models focus on the properties of monolayers as a bulk. However, as Devany et al. showed in simulations and experiments that absolute changes of the cell shape can vary greatly and could thus be inconclusive, depending on the experimental situation.^[62]^ Importantly, Saraswathibhatla and Notbohm found a correlation between cell density, shape, and motility.^[63]^ While we only observe small changes in cell shape, we do observe a similar impact of cell density.

In addition, most studies identifying the cell shape as the predictive parameter for cell motility worked with other cell types and on longer times scales. Typically, fully polarized cells, such as airways smooth muscle cells cultivated for several days and up to weeks, were used, whereas our MDCK II cells only had about 28 h to grow to full (over-)confluence.^[21,26,64]^ Studies working with (ZO protein-inhibited) MDCK or MCF10A lines also cultivated the cells much longer,^[2,65]^ which in conjunction with the observed uncontrolled proliferation could explain the density-related discrepancies. Furthermore, related studies investigated the motion of confluent cell layers, whereas we focused on freely migrating epithelia.^[21,26,31,63–65]^

## 4. Conclusion

We showed that ZO proteins are not only crucial for barrier function but also required for efficient collective cell migration of epithelial monolayers. Our results draw the following picture of the impact of ZO1 and 2 protein loss: Due to missing mechanical feedback from ZO1/2, a thick actomyosin ring builds at the cell periphery that leads to strong contraction of individual cells, constricting the apical cell cortex and leading to in- or outward bulging. In order to keep the adhesion to the substrate not all cells can adopt this morphology and competition between contractility and adhesion emerges. As a consequence, two subpopulations of cell phenotypes arise in ZO1 and 2 depleted cells after a few hours of migration: 1) Small contractile cells (‘winner’) with an apically bulged-out and softened cortex and 2) large, flat cells (‘loser’) with an elevated prestress. The larger cells respond to the mechanical stimulus from the highly contractile neighbors by increased proliferation, leading to more immobile ‘winner’ cell and eventually to crowding that slows down migration of the cell sheet. We conclude that functioning tight junctions are necessary for tension homeostasis to maintain fluidity of epithelial monolayers and thereby guarantee for fast and coherent cell migration.

## 5. Material and Methods

### Cell culture

Madin-Darby Canine Kidney cells (strain II, MDCK II; European Collection of Authenticated Cell Cultures, Salisbury, UK) were cultured in minimum essential medium (MEM; Life Technologies, Paisley, UK) containing Earle’s salts, 2 mM GlutaMAX^TM^ (ThermoFisher Scientific, Waltham, Massachusetts, USA), 2.2 g/L NaHCO_3_, and 10% fetal bovine serum (FCS; BioWest, Nuaillé, France), called M10F^-^ in the following, at 37°C and 5% CO_2_ in a humidified incubator. The cells were passaged before reaching confluence two to three times per week using phosphate buffered saline pH 7.4 (PBS^-^; Biochrom, Berlin, Germany) containing trypsin/EDTA (0.25%/0.02% w/v; BioWest/Biochrom).

### Genetic modification of ZO proteins

ZO knockdowns were effected as described in Beutel et al.^[45]^ To knockdown ZO1 and ZO2 in MDCK II cells, frame-shift mutations were introduced at the N-termini by CRISPR/Cas9. The following RNA guides (gRNA) were used for ZO1: ACACACAGTGACGCTTCACA and ZO2: GTACACTGTGACCCTACAAA. Selected DNA oligos and their trans-encoded RNA (TRCR) were purchased from Integrated DNA Technologies. Each gRNA was annealed for 1h at room temperature with its TRCR. To finally generate the riboprotein complex, the gRNA/TRCR complex was incubated with homemade purified Cas9. Electroporation of each complex was performed in 300,000 cells (Invitrogen NEON electroporation machine and kit, 2 pulses, 20 ns, 1200 V). Single cells were sorted after 48 h by FACS (fluorescence activated cell sorting) and grown clonally. The genomic sequence of the genes of interests were sequenced and only clones carrying homozygous frame-shifts leading to an early stop codon were kept. To generate a combined ZO1/ZO2 knockdown KD line, we first created a ZO1 knockout and then we targeted ZO2. The ZO1 KO clone was mutant for two alleles, both alleles have a 1 bp insertion in the guide region (ACACACAGTGACGCTTC-1 bp insertion-ACAGGG) leading to an early stop of translation. The ZO2 KD has 5 bp deletion at the end of the guide region (GTACACTGTGACCCTACA-5 bp deletion-GG) leading to an early stop. Immunostaining and western-blot analysis showed that ZO1 and ZO2 presented a residual expression level of the full-length protein equal to 2-3% of the WT line expression level (Beutel et al Fig S5).

### Generation of MDCK II WT-GFP cells

Clones of MDCKII expressing GFP-myosin-2-A were created by transfecting cells with pTRA-GFP-NMCH II-A plasmid (Addgene plasmid # 10844). Stable expressing clones were selected via Neomycin resistance (G418). After selection, the cell pool was sorted by FACS to enrich for cells expressing GFP at a moderate level.

### Cell migration experiments

For migration experiments Petri dishes with a culture-insert (Culture-Insert 2 Well in µ-Dish 35 mm, ibiTreat #1.5 polymer cover slip; ibidi, Martinsried, Germany) were used. Cells were seeded at 4·10^5^ cells in 1 mL M10F^-^ on the outside of the insert and grown to (over-) confluence for 28 h (± 1.5 h). WT-GFP/dKD co-cultures were trypsinized and mixed well before being seeded simultaneously at 2·10^5^ each (1:1 ratio) and grown as described above. Upon visual inspection, the insert was removed, the cells were rinsed with M10F^-^, supplied with sufficient M10F^-^ (2-3 mL), and placed into the incubation system of an inverted optical microscope (BZ-X810; Keyence, Neu-Isenburg, Germany) equipped with a 10X phase contrast objective (Nikon CFI60 Series; Keyence). The temperature was calibrated to be 37°C at the cell sample using a local temperature measurement instrument (Testo 735; Testo, Lenzkirch, Germany), a partial CO_2_ pressure of 5% was chosen, and sufficient humidity was ensured by injecting distilled water into the incubator appliance. Phase contrast frames were recorded at 1 frame/2.5 min, 14 bit, 25% illumination power, typical exposure times of about 1/25 s, and without zoom, gain, or binning. Focus tracking was applied and three vertical slices were chosen in a range of 5 µm to avoid drift effects. The cell edge was carefully aligned vertically and set to be at a similar position for all experiments. Typically, migration was observed overnight for 20-30 h.

### Mitomycin C treatment

Mitomycin C (MitoC; Sigma-Aldrich, Steinheim, Germany) was dissolved in water to reach 500 µg mL^−1^ and stored in aliquots of 150 µL.

Cell seeding was performed as described above. The samples were rinsed once and then incubated with M10F^-^ containing 10 µg mL^−1^ of Mitomycin C at 37°C and 5% CO2 for 1 h. Then, the insert was removed after about 28 h growth time (± 1.5 h). To remove any extruded cells and, most importantly, to prevent the cytotoxic effects of Mytomycin C occurring after 12 h of exposure,^[18]^ samples were rinsed with 1 mL M10F^-^ three times, before the dishes were filled with 2-3 mL M10F^-^ and then imaged as described above.

### ROCK inhibition by Y27632 treatment

Y27632 (*”*InSolution” Y27632; Sigma-Aldrich) was diluted in M10F^-^ to the desired concentration, cells were rinsed once and then incubated in the Y27632-containing medium. In the case of continuous treatments, 30 min of incubation was allowed before insert removal, cells were rinsed again and 2-3 mL Y27632-containing medium was added.

### Experiments with non-migrating monolayers

5 10^5^ cells were seeded in 4mL M10F^-^ and placed immediately on the same microscope as above and the same conditions as for the migration experiments were used but without an insert. Four areas per sample were imaged every hour with the same settings as above. Two WT samples, one KO and one dKD sample were recorded. Analysis was performed as described below.

### Migration data analysis

First, migration phase contrast movies were down-sampled to 1 frame/7.5 min to ensure good PIV (particle image velocimetry) quality. Velocity vector maps were obtained using the Matlab (MathWorks, Natick, USA) -based PIV tool AVeMap from Deforet et al.^[35]^ A window size of 32 x 32 pixels corresponding to 24.16 µm x 24.16 µm with an overlap of 0.5 was used, yielding a vector mesh size of 16 pixels (12.08 µm). The first row width was set to 12.08 µm and typical mask parameters were 0.60-0.75. The default filters of 1.1 signal-to-noise ratio, 0.3 peak height, and 4 global filtering were used. A PIV quality of > 0.8 was achieved for all data and exemplarily checked by visual inspection. The order parameter was defined as cos *α*, where *α* is the angle between the local velocity vector and the normal to the average migration direction. The add-on AVeMap+ was used to analyze the data with respect to the distance from the migration edge. Note, the first two to three data points are not shown due to a known edge-induced artefact.^[66]^

Vector fields were further analyzed using home-written Python scripts. Before correlation functions were calculated, the leader cell fingers were cut from the vector fields to yield rectangular input data for the spatial correlation and to avoid edge-induced artifacts.

The correlation function was calculated for each time point individually as the 2D spatial autocorrelation *AC* of the velocity vector field using the Scipy function signal.correlate2d and according to Petitjean et al.:^[32,67]^

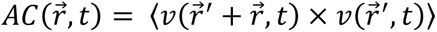

With the deviation of the *y*-component (perpendicular to the migration direction) *ν* = *ν_i_* − 〈*ν*〉, which is corrected by the offset 〈*ν*〉, of the vector 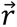 at time point *t*. The brackets denote averaging over all vectors. Additionally, the *AC* is normalized by its maximum, so that it starts from one.

To gain a one-dimensional function, the 2D correlation function was then radially averaged in space. The correlation function was finally averaged for each migration movie over time.

The correlation length was defined as the integral over the weighted spatial correlation function *AC*(*r*):

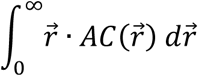

To exclude any anti-correlation artifacts (*AC* < 0) at large distances, we only integrated up to the x-intercept for all analyses.

The amount of leader cells was determined from the phase contrast movies manually. Leader cells were defined by their position at a protrusion in the front of the leading edge, an increased cell size compared with bulk cells, and a lamellipodium towards the empty space.

### Automated cell segmentation

The deep learning-based cell segmentation algorithm Cellpose (Stringer et al.^[38]^) was used to extract a mask and an outline for each individual cell body in an image. The model type was set to *cyto* and the grayscale phase contrast images were used as input. Before segmentation, the image contrast was auto-corrected using Fiji to facilitate optimal cell recognition.^[68]^ In order to accurately capture all cells in the layer, the flow and cell probability thresholds were set to 1 and −6, respectively, for the phase contrast monolayer images. The parameters for other analyses were set as follows: Single cell phase contrast: 0.7 and −2. Mixed monolayer GFP-fluorescence-channel: 0.95 and −5. Confocal YAP- and corresponding DAPI-images: 0.4 and 0 (Cellpose default values). Confocal actin-images: 0.7 and −2. Diameters were detected automatically, except for confocal images, where they were pre-adjusted by visual inspection.

We found these parameters to be optimal for our images, because smaller (or larger, respectively) values resulted in missed cells. No novel model training was necessary. The input diameter was estimated automatically for every image individually by the software.

### Cell area, position, and aspect ratio calculation and processing

For every segmented image, the masks array and the outlines array were extracted from the returned segmentation dictionary. The arrays were normalized, so that ones specified cell bodies (or cell outlines) and zeros empty space, respectively. The outlines were subtracted from the masks to prevent overlap of cells. The resulting array was converted into the data type uint8 and scaled up to a value of 255. The array was then subjected to a threshold at a value of 127 and binarized using the image processing library OpenCV.^[69]^ The arrays were then transposed into vectors of coordinates specifying the outer contour of each cell using the function *findContours* of OpenCV.^[69]^ Only outer contours were extracted and the Teh-Chin chain approximation algorithm was applied to save memory.^[70]^ On the basis of the extracted vectors, the area of each cell was computed using the function contourArea of OpenCV. The moments function was used to determine the center of each cell, yielding the positions later used by Trackpy. Cell density was calculated dividing the number of segmented cells by the area occupied by the monolayer (either the mask obtained from AVeMap or the whole field of view).

To determine cell aspect ratios (length/width), two approaches were utilized to define the front-rear (anterior-posterior) axis for each individual cell. First, the fitEllipse function of OpenCV was used for every given set of coordinates to compute and fit an ellipse to the 2D points. Since this function works by fitting the coordinates in a least-squares approach, it was found that the algorithm seemed to be biased towards high aspect ratios for some cell shapes. Therefore, the function minAreaRect was used to verify the results by calculating a rotated minimum-area rectangle enclosing the respective set of coordinates. This procedure, however, seemed to be biased towards low aspect ratios for the aforementioned cell shapes. Accordingly, we computed the aspect ratio with both algorithms independently and then used the mean for every cell in each image individually. The validity of this approach was verified by visual inspection of overlaid input and output images.

### Cell tracking and analysis

Single cell tracking was performed with the cell positions calculated before by the OpenCV moments function (*vide supra*). Trackpy was used to link the cell positions, yielding individual tracks.^[71,72]^ The link function was used with a memory of 4 frames and 4.6 or 5.7 µm^2^ (8 or 10 pixels) as maximal displacement (10 frames and 20 pixels for single cell migration). The resulting trajectories were filtered, so that only the ones that persisted for at least 5 frames were kept, to avoid spurious trajectories. No drift correction was necessary. The temporal resolution was 1/7.5 min for all monolayer data and 1/2.5 min for the single cells. Cell velocities were calculated for each cell by averaging over all time steps, which automatically normalizes for the different frame rates.

Mean-square-displacements (*MSD*s) were calculated using the ensemble *MSD* function of Trackpy as:

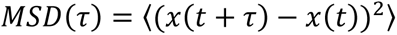

The brackets denote averaging over time and over all cells. Before calculation and fitting of the *MSD*s, the trajectories were filtered by discrete bins of 100 µm^2^ cell area (see Figure 2 and 3) or 0.25 aspect ratio (see Figure S1 and S2). *MSD*s were fitted by a power law of the form *MSD*(*τ*) = *a τ^n^* with a power law exponent *n* and an offset *a* using a linear regression in logarithmic space implemented in Trackpy.

### AFM-based force spectroscopy

Force spectroscopic indentation measurements were carried out with a NanoWizard 4 (JPK Instruments, Berlin, Germany) mounted on an inverted microscope (IX 81; Olympus, Tokyo, Japan) using silicon nitride cantilevers with a nominal spring constant of 0.01 N m^−1^ (MLCT C; Bruker AFM Probes, Camarillo, USA). Before an experiment, cantilevers were rinsed with isopropanol and PBS^-^ as well as incubated with FITC-conjugated Concanavalin A solution (2.5 mg mL^−1^ in PBS^-^; Sigma-Aldrich) for 1 h.

The sensitivity of the AFM was determined by recording force curves in the empty space without cells and the exact spring constant of each cantilever was determined by the thermal noise method.^[73]^ Approximately 20 h after removing the insert (*vide supra*), cells were rinsed three times with M10F^-^ containing 0.2 mg mL^−1^ Penicillin (Biochrom), 0.2 mg mL^−1^ Streptomycin (Biochrom), and 15 mM HEPES (M10F^+^; BioWest).

For the measurements, samples were mounted on the AFM stage, 2.5 mL M10F^+^ was supplied, and the heater (JPK Instruments) was set to 37°C. The cells were indented at a constant speed of 2 µm s^−1^ to maximum force of 1 nN. After a dwell time of 0.5 s at constant height the indenter was retracted at the same speed. Force maps of 25 pixels x 25 pixels in an area of 50 µm x 50 µm were recorded by lateral scanning across the sample recording one force indentation cycle at each pixel. Additionally, five consecutive force curves in the center of individual cells in the monolayer were acquired using the same parameters.

### Force curve analysis and mechanical model

Generally, force-relaxation curves were recorded as detailed previously.^[46]^ After indentation of the center of the cell with a velocity of 2 µm s^−1^ to avoid artefacts from hydrodynamic drag acting on the cantilever, we switched off the constant force feedback loop and kept the system at constant height. During this time the decrease of cantilever deflection is monitored as a function of time (for 0.5 s). We used the same MLCT-C cantilevers as for imaging (*vide infra*). The curves were modeled using a theory introduced recently.^[46,74]^ Briefly, the surfaces of the confluent MDCK II cells are described as capped cylinders. The average geometry as derived from AFM imaging was employed to describe the apical cap of the deformed cells in terms of contact angle and radius of the basis. Generally, we consider the cell as a liquid-filled object surrounded by an isotropic viscoelastic shell deformed at constant volume. The force *F* acting on the apex of the cell is given by:

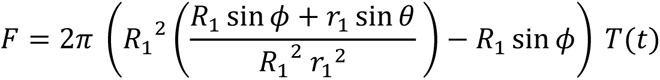

with *R*_1_, the radius at the base of the spherical cap and *ϕ* the contact angle in response to deformation. *r*_1_ is the contact radius with the conical indenter, 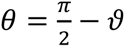 with *ϑ*, the cone half angle.

Viscoelasticity of the shell enters the tension term *T*(*t*) through a time dependent area compressibility modulus *K_A_* = *K_A_*^0^(*t*/*t*_0_)^−*β*^. Now we need to solve a set of nonlinear equations for the shape of the deformed cell to fulfill force balances and the constant volume boundary condition. The resulting shapes are minimal surfaces to minimize the stretching energy. Membrane tension *T*_t_ was calculated from the tether rupture force *F*_t_ at the end of the retraction curve via 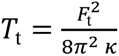 with the bending modulus *κ* = 2.7 · 10^−19^ J.^[75–77]^

Analysis was performed using self-written Python and Matlab scripts in combination with the JPK SPM Data Processing (JPK Instruments / Bruker) software. The baseline was corrected by a linear fit before contact. The contact point was determined individually using the JPK SPM Data Processing. Tether forces were acquired with the same software.

### AFM imaging

Approximately 20 h after removing the insert (*vide supra*), cells were rinsed three times with PBS containing 0.1 g L^−1^ Mg^2+^ and 0.133 g L^−1^ Ca^2+^ (PBS^++^; Sigma-Aldrich) and incubated with glutaraldehyde solution (2.5% (v/v) in PBS^++^) for 20 min. PBS^++^ was used instead of PBS without magnesium and calcium ions, because the dKD and KO cells were more prone to dissolution of ion-dependent adhesions due to the missing diffusion barrier function. Before imaging, the samples were rinsed again three times to remove residual GDA. Cell imaging was performed using a NanoWizard III (JPK Instruments) mounted on an inverted optical microscope (IX 81; Olympus) to enable additional visual inspection via phase contrast. Imaging was carried out as described in Brückner et al.^[49]^ in contact mode using MLCT C cantilevers (Bruker AFM Probes) in PBS^++^ with typical line scan rates of about 0.3 Hz and typical forces of 0.1 nN. Height and error images were obtained using the JPK SPM Data Processing software provided by the manufacturer.

### Cell labeling and fluorescence microscopy

Prior to cell labeling, cells were fixed by incubation with paraformaldehyde/glutaraldehyde solution (4% (w/v)/0.1% (w/v) in PBS^++^; Science Services, Munich, Germany/Sigma-Aldrich) for 20 min. To permeabilize the cellular plasma membrane, samples were incubated with 0.1% (v/v) Triton X-100 in PBS^-^ for 5 min. After three rinsing steps with 1 mL PBS^++^ each, to block unspecific binding sites, cells were incubated with blocking/dilution buffer (PBS^-^ containing 2% (w/v) bovine serum albumin and 0.1% (v/v) Tween20) for 30 min.

For ZO1 staining, a fluorophore-conjugated primary antibody (mouse ZO1-1A12 IgG1 AlexaFluor 488; Invitrogen, ThermoFisher Scientific, Waltham, Massachusetts, USA) was diluted with blocking/dilution buffer to a concentration of 5 µg mL^−1^ and cells were incubated for 1 h. For all other proteins, the following primary antibodies were diluted in blocking/dilution buffer.

### ZO2

1 µg mL^−1^ Clone 3E8D9 mouse IgG1; Invitrogen. Phospho-Myosin: 1:200 light chain 2 (Ser 19) rabbit IgG1; Cell Signaling Technology, Danvers, Massachusetts, USA. *E-Cadherin*: 1:50 Clone 36 mouse IgG1; BD Biosciences, Heidelberg, Germany. *ß-catenin*: 5 µg mL^−1^ mouse IgG1; BD Biosciences. *Occludin*: 6.5 µg mL^−1^ EPR20992 rabbit IgG; Abcam, Cambridge, UK. Claudin 1: 11.6 µg mL^−1^ rabbit IgG; Abcam. *YAP*: 1:100 (5-10 µg mL^−1^) Anti-YAP1 rabbit, SAB2108066; Sigma-Aldrich.

After the primary antibody, cells were rinsed briefly with PBS^++^, and then washed with PBS^++^, with 0.1% (v/v) Triton X-100 in PBS^-^ and again with PBS^++^ for 5 min each on a shaker plate (75 rpm). The secondary antibody (AlexaFluor 488- or AlexaFluor 546-conjugated goat anti-mouse or anti-rabbit IgG; Life Technologies, Carlsbad, USA) was diluted with blocking/dilution buffer to a concentration of 5 µg mL^−1^. The cells were incubated for 1 h. Actin labeling was performed using AlexaFluor 488- or AlexaFluor 546-phalloidin (Invitrogen), diluted together with the secondary antibody in blocking/dilution buffer to a concentration of 165 nM. Incubation time: 45 min. Following the secondary antibody, samples were washed with PBS^++^ for 5 min each on a shaker plate (75 rpm). Nucleus staining was performed by incubation with Hoechst 33342 (Invitrogen), diluted to 1 µg mL^−1^, for 15 min. For imaging, samples were rinsed three times with PBS^++^ and kept in PBS^++^. All labeling steps were performed at room temperature.

A confocal laser scanning microscope (FluoView1200; Olympus, Tokyo, Japan), equipped with a 60x oil immersion objective (*NA* = 1.25), was used for fluorescence imaging. Image processing, brightness adjustment, and analysis was performed using Fiji.^[68]^

### Nucleus/cytoplasm-localization quantification of YAP

As described above, Cellpose was used to segment the cytoplasm and nucleus for each cell in a confocal image of the central plane. Segments were used as masks to extract the respective intensities (YAP in the cytoplasm and nucleus) from the original image. For each cell, the mean intensity in the nucleus was divided by the mean intensity in the nucleus to, yielding a ratio between nucleus and cytoplasm, i.e. the relative nucleus localization of YAP. Values above 1, which resulted from false segmentation, were excluded from the results.

### Cell volume analysis

The base cell area was determined using Cellpose and OpenCV at the basal side of the cell from confocal actin-images. The cell height was determined visually from side views in Fiji. Cell volume was calculated by multiplying area and height. Lastly, the theoretical isotropic expansion was calculated for a cylinder, considering proportionally even changes in radius and height.

### Statistical analyses

The data were tested for normality using the Shapiro-Wilk test. Because for none of the PIV-based data (Figure 1, and Figure 3B) the null hypothesis of a normal distribution was rejected (at the *p* < 0.05 level), significance was tested using Welch’s t-test. The Mann-Whitney U test was applied to the rest of the data to accommodate non-normality. All statistical analyses were performed in Python.

A *p*-value of < 0.05 was considered significant and denoted by one asterisk (*), *p* < 0.01 and *p* < 0.001 we indicated by two (**) and three (***) asterisks, respectively.

## Supporting Information

Supporting Information is available at the end of this document.

## Acknowledgements

Funding by the DFG through grants SPP1782 and DFG JA963 is gratefully acknowledged. We thank Jonathan F.E. Bodenschatz and Justus Schünemann for technical assistance and discussions. We would also like to acknowledge Filip Savić, for his help with the PIV analyses and Burkhard Geil for the helpful discussions about the segmentation and tracking. M.S., H.P., M.F., J.G., and A.R. executed measurements and performed analyses. M.S. designed and planned the experiments. R.M. and A.H. carried out the genetic modifications. T.O., A.H., and A.J. designed and supervised the research. M.S. and A.J. wrote the manuscript. All authors helped with discussions and proofreading.

---

In this article it is shown that tight junction ZO proteins maintain epithelial cell sheet fluidity and thereby ensure efficient and coherent migration. Cells lacking these proteins loose viscoelastic tissue integrity due to actomyosin remodeling and increase proliferation, which induces cellular crowding. Particularly, upon ZO protein loss small cells at high cell densities eventually impair migration and cause jamming.

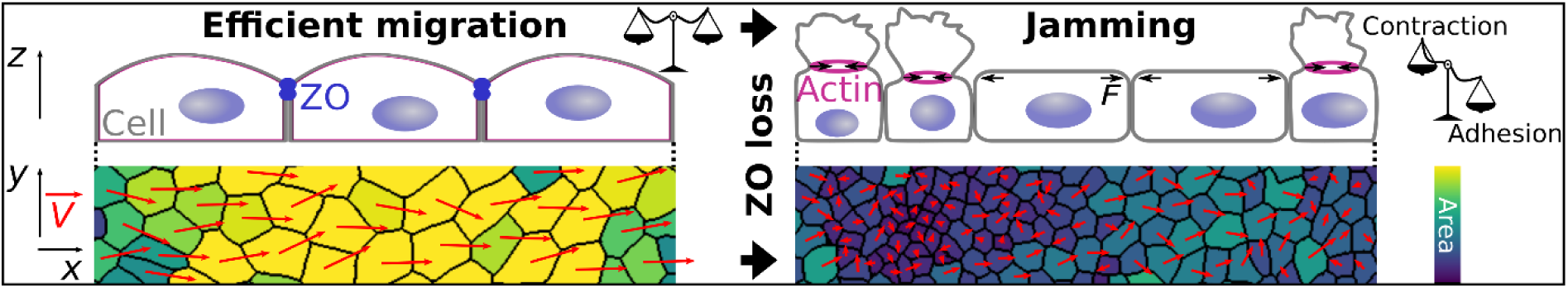
ToC figure

## Supporting Information

**Figure S1.**
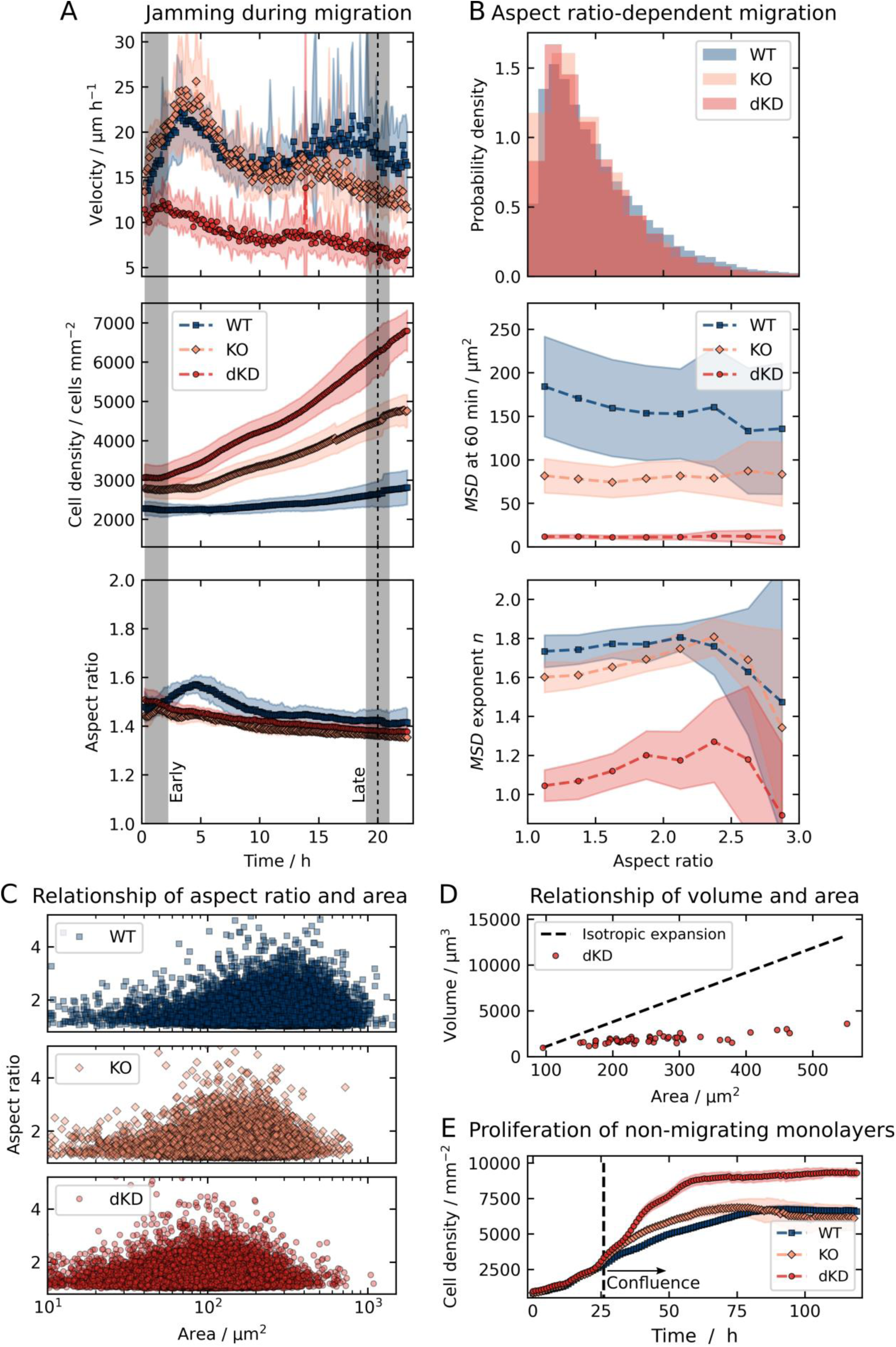
Temporal evolution of cell density, velocity, and aspect ratio as well as aspect ratio-dependent motility of all three untreated MDCK II cell lines. A) Cell crowding and jamming during migration as quantified by the velocity, cell density, and aspect ratio over time. The gray shades at 0.5-2.5 h (‘early’) and 19-21 h (‘late’) correspond to the time windows of the cell velocity and *MSD* analyses. B) Aspect ratio distribution and aspect ratio-dependent MSD parameters. C) The aspect ratio showed a high variance but no co-variation with the area. D) Relationship between the increase in cell volume with respect to the cell area in dKD cells determined from 3D-confocal actin stacks. For comparison, theoretical isotropic expansion is shown as a dashed line. E) Additional proliferation experiment immediately after seeding of cells without insert. The dashed line indicates reaching of confluence, corresponding to the beginning of our typical migration experiments (0 h in all other figures). Means and standard deviations are shown. The aspect ratio in A is the median for each experiment and then averaged over all experiments.

**Figure S2.**
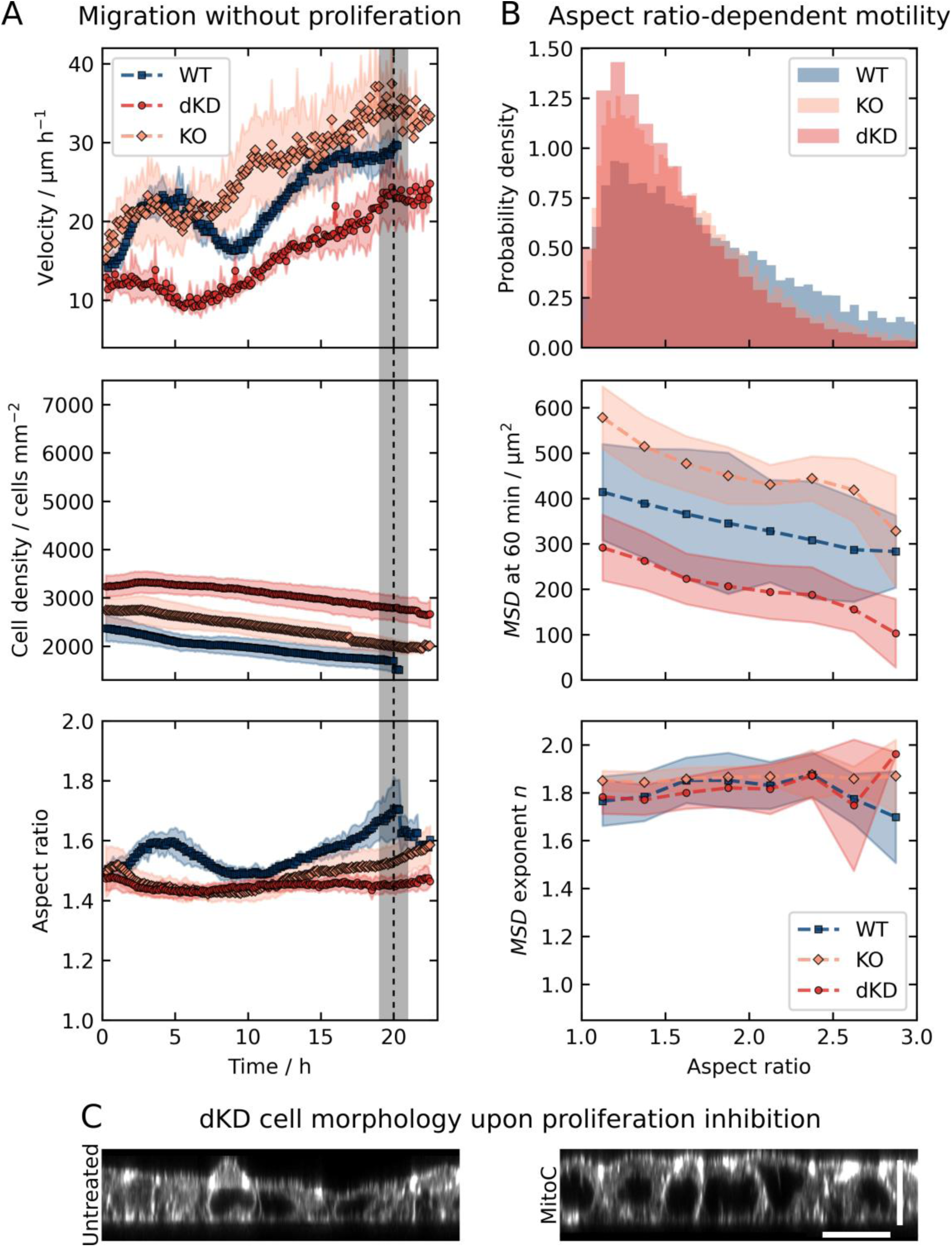
Temporal evolution of cell density, velocity, and aspect ratio as well as aspect ratio-dependent motility of all three MDCK II cell lines upon proliferation inhibition by Mitomycin C. A) Cell crowding and jamming were prevented by proliferation inhibiton during migration as quantified by the velocity, cell density, and aspect ratio over time. The gray shade at 19-21 h corresponds to the time window of the MSD analyses. B) Aspect ratio distribution and aspect ratio-dependent MSD parameters. Means and standard deviations are shown. The aspect ratio in A is the median for each experiment and then averaged over all experiments. C) Side-view of untreated and Mitomycin C treated dKD cells from 3D confocal actin stacks. Scale bars: 10 µm (z), 20 µm (x).

**Figure S3.**
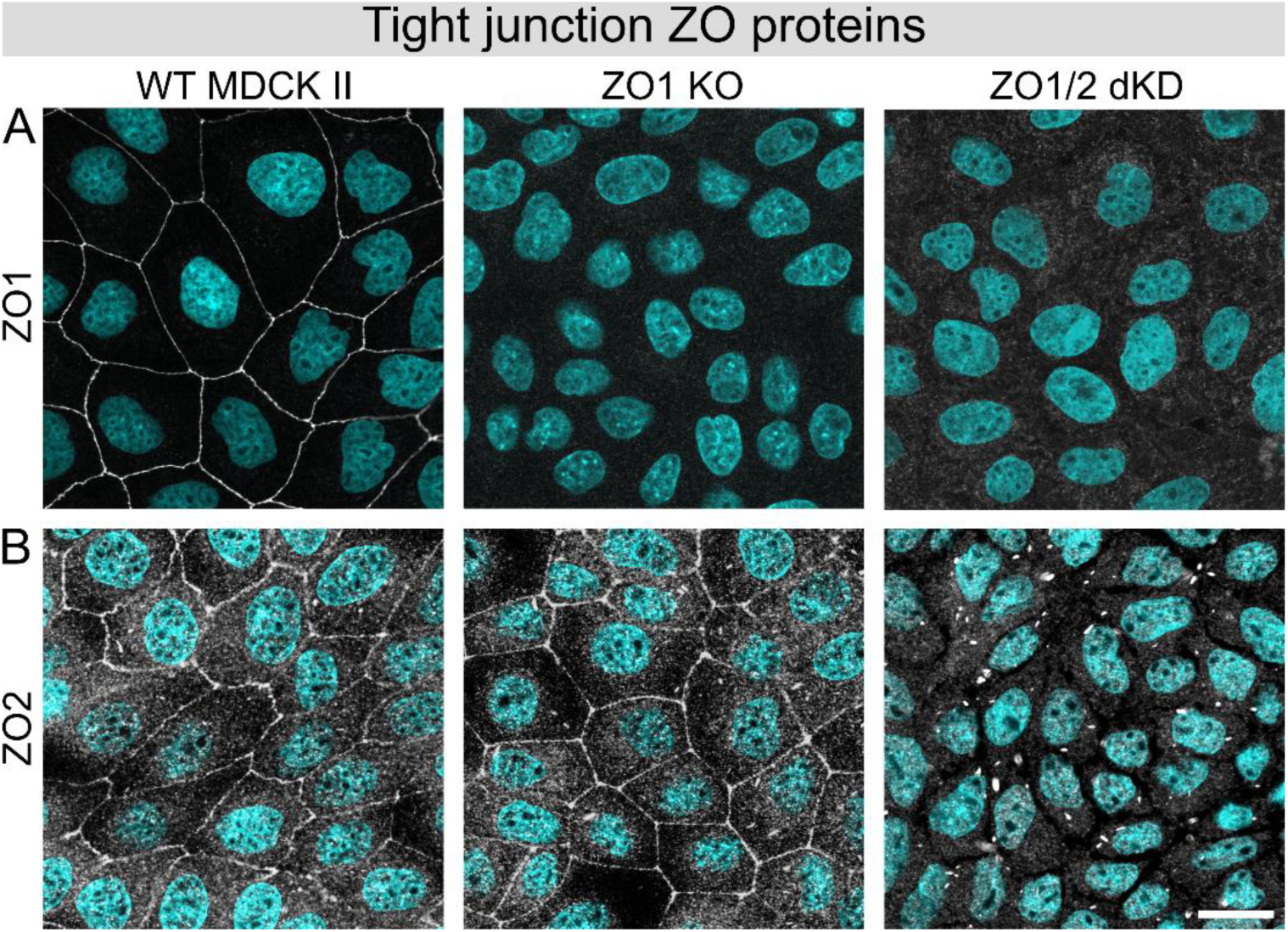
Immunofluorescence measurements confirming successful ZO protein knockout/-down. A) ZO1 antibody-based staining of all three MDCK II cell lines. B) ZO2 antibody-based staining of all three MDCK II lines. Nuclei are shown in cyan. Scale bar: 20 µm.

**Figure S4.**
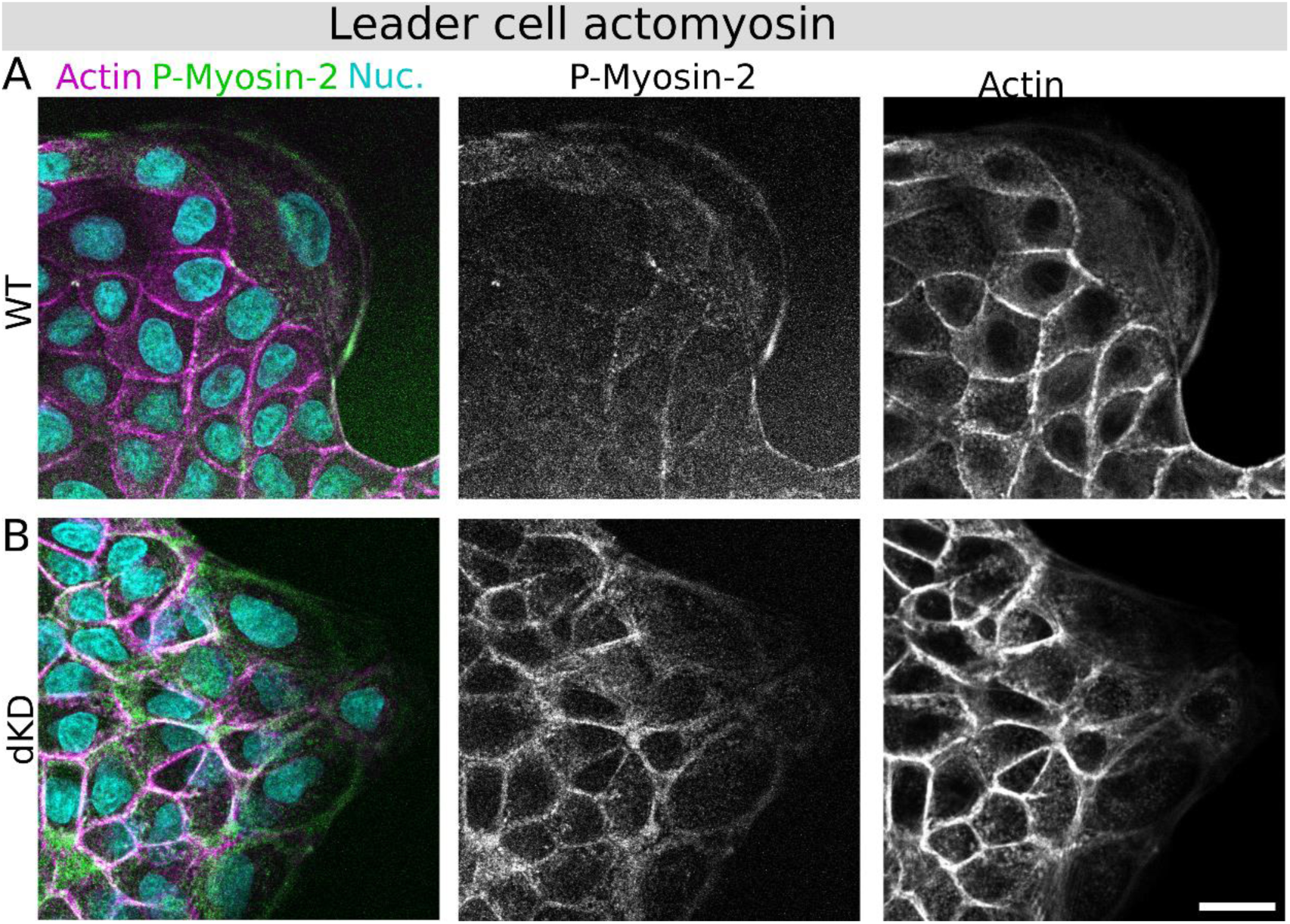
Actomyosin architecture remodeling for leader cells at the migration front of migrating WT and dKD cell layers. Phosphorylated Myosin-2 (P-Myosin-2; green), actin (magenta) and nuclei co-staining of MDCK II WT (A) and dKD (B) cell lines with corresponding gray-scale images of P-Myosin-2 and actin. Scale bar: 20 µm.

**Figure S5.**
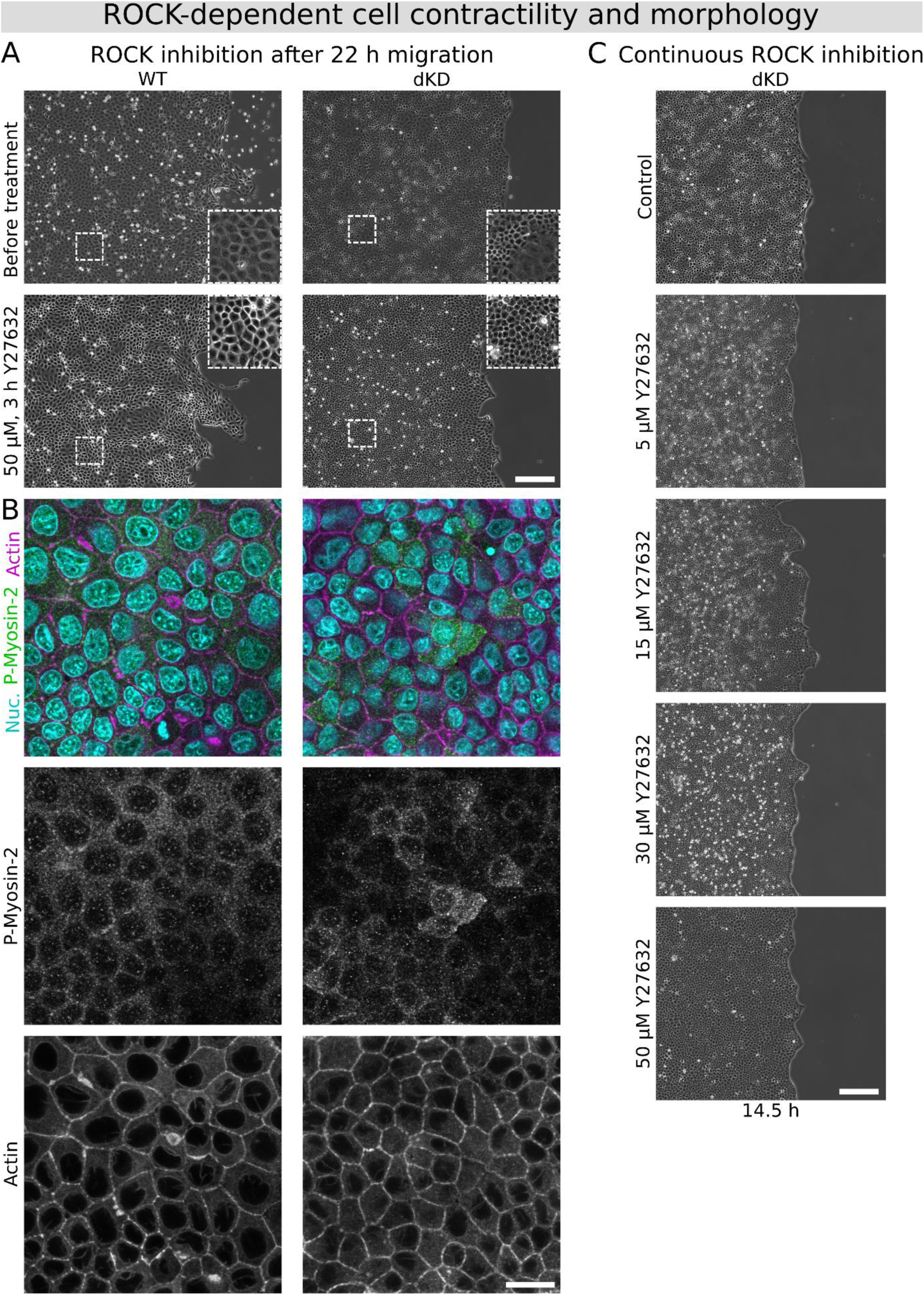
MDCKII WT and dKD cell lines show altered migration and actomyosin architecture remodeling upon inhibition of ROCK with Y27632. A) Inhibition of ROCK by Y27632 after 22 h of migration in WT and dKD cells. Top shows cell monolayers before treatment and bottom the same layers after 3 h of migration in the presence of 50 µM Y27632. Scale bar: 200 µm B) Confocal actomyosin images of the same MDCKII WT and dKD layers from A after Y27632 treatment after 22 h of migration with corresponding gray-scale images of P-Myosin-2 and actin. Scale bar: 20 µm. C) Migration of dKD cells for 14.5 h in the presence of increasing concentrations of Y27632 (starting continuously from 30 min before insert removal). Scale bar: 200 µm.

**Figure S6.**
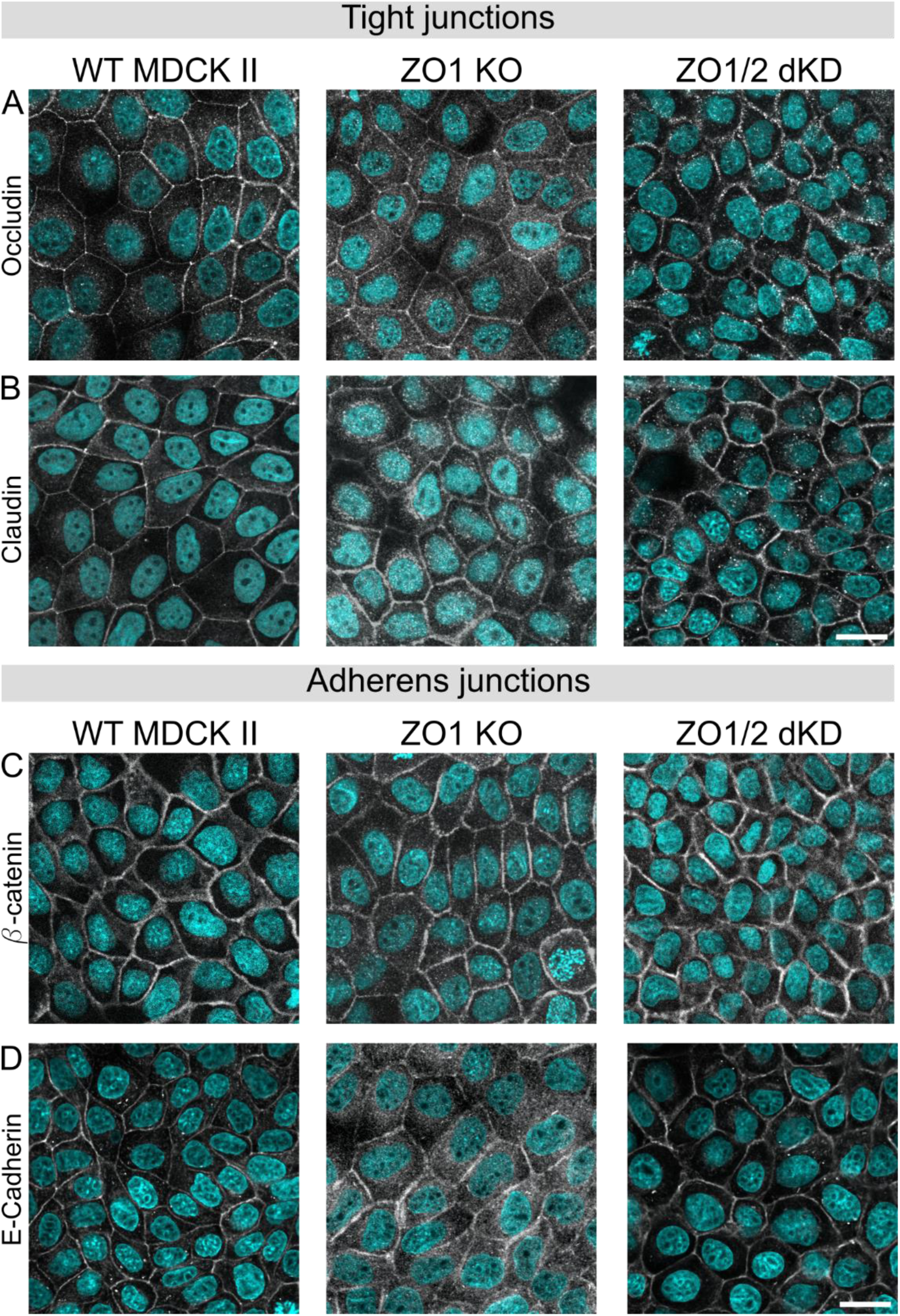
Immunofluorescence of tight junction transmembrane proteins and adherens junction proteins. A) Occludin. B) Claudin 1. C) β-catenin. D) E-Cadherin. Nuclei are shown in cyan. Scale bars: 20 µm.

**Figure S7.**
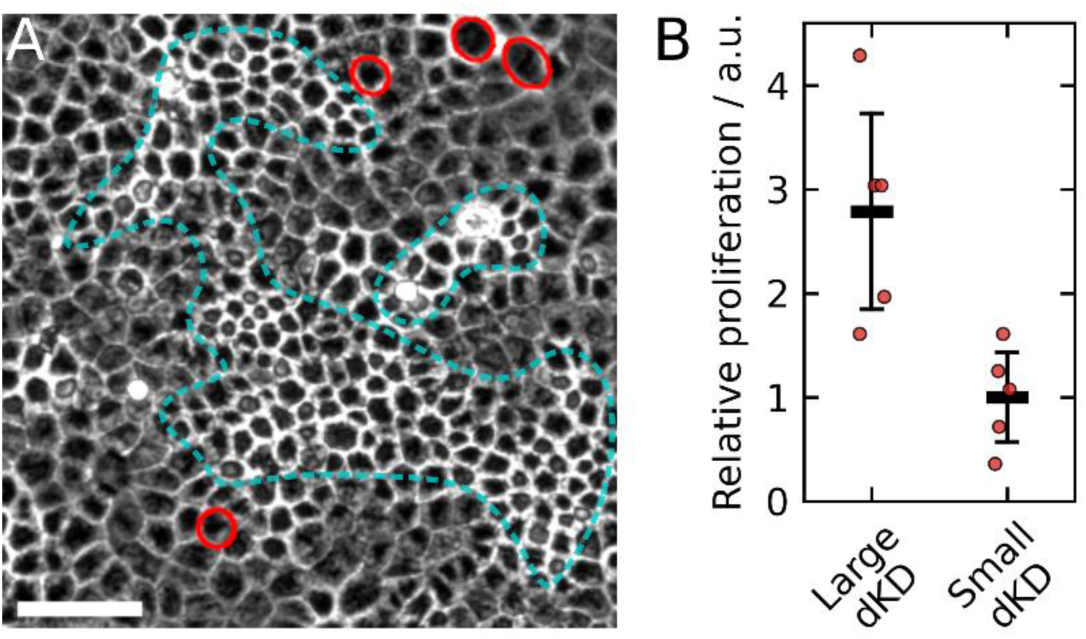
In the dKD monolayers, more of the large and stretched cells were observed to proliferate than the small and contractile cells. A) Example of dKD cells during migration with proliferation events indicated by red circles and patches of small cells highlighted in cyan. Scale bar: 50 µm. B) Relative proliferation of five example regions from A, normalized by the average number of small cell proliferation events. Proliferation events were counted and attributed by hand and the examples were chosen, so that approximately the same amount of large and small cells was present. These data were collected in the same time window as the MSD analysis, i.e., between 19 h and 21 h.

